# T cell receptor-centric perspective to multimodal single-cell data analysis

**DOI:** 10.1101/2023.09.27.559702

**Authors:** Kerry A. Mullan, My Ha, Sebastiaan Valkiers, Nicky de Vrij, Benson Ogunjimi, Kris Laukens, Pieter Meysman

## Abstract

The T-cell receptor (TCR) carries critical information regarding T-cell functionality. The TCR, despite its importance, is underutilized in single cell transcriptomics, with gene expression (GEx) features solely driving current analysis strategies. Here, we argue for a switch to a TCR-first approach, which would uncover unprecedented insights into T cell and TCR repertoire mechanics. To this end, we curated a large T-cell atlas from 12 prominent human studies, containing in total 500,000 T cells spanning multiple diseases, including melanoma, head-and-neck cancer, T-cell cancer, and lung transplantation. Herein, we identified severe limitations in cell-type annotation using unsupervised approaches and propose a more robust standard using a semi-supervised method or the TCR arrangement. We then showcase the utility of a TCR-first approach through application of the novel STEGO.R tool for the successful identification of hyperexpanded clones to reveal treatment-specific changes. Additionally, a meta-analysis based on neighbor enrichment revealed previously unknown public T-cell clusters with potential antigen-specific properties as well as highlighting additional common TCR arrangements. Therefore, this paradigm shift to a TCR-first with STEGO.R highlights T-cell features often overlooked by conventional GEx-focused methods, and enabled identification of T cell features that have the potential for improvements in immunotherapy and diagnostics.

**One Sentence Summary:** Revamping the interrogation strategies for single-cell data to be centered on T cell receptor (TCR) rather than the generic gene expression improved the capacity to find relevant disease specific TCR.

**Key Points:**

- The TCR-first approach captures dynamic T cell features, even within a clonal population.
- A novel ∼500,000 T-cell atlas to enhance single cell analysis, especially for restricted populations.
- Novel STEGO.R program and pipeline allows for consistent and reproducible interrogating of scTCR-seq with GEx.

## Introduction

T cells are specialized cells of the adaptive immune system positioned to recognize various infected or otherwise aberrant cells. This recognition is driven by the T-cell receptor (TCR) expressed on the cellular surface. Their importance for the immune response, and various underlying disease states, have driven recent innovations in the domain of single-cell sequencing. We can now capture a multimodal view of a single T cell that includes the transcription gene expression (GEx), as well as the hypervariable TCR(*1*). This detailed perspective allows distinction of the T cell population into its phenotypes with the cognate TCRs. The broadest phenotypic distinction is between CD4+ T cells, which recognize the class II major histocompatibility complex (MHC) loaded with extracellular peptide sources, and CD8+ T cells that recognize the class I MHC loaded with intracellular peptides. CD4+ T cells exhibit diverse functional phenotypes, categorized into T helper (Th) subtypes such as Th1, Th2, Th9, Th17, follicular T cells (Tfh), and regulatory T cells (Tregs). T cells also display a variety of memory states and can transition from activation to senescence or exhaustion. In addition to their expression markers, T cells are divided into two principal lineages based on the T cell receptor (TCR), namely alpha-beta (αβ) or gamma-delta (γδ). Each of these four TCR chains is made up of a Variable (V), Diversity (D) and Junction (J) genes, with alpha/gamma having VJ and beta/delta VDJ arrangements. During development a T cell will acquire a quasi-random recombination of V(D)J genes, which results in a hypervariable sequence in the complementarity determining region 3 (CDR3), the primary driver of variable antigen specificity. Recent methodological advances have enhanced the analysis of bulk TCR repertoire data, with novel tools based on identifying antigen-specific TCR groups (e.g., ClusTCR(*2*), tcrdist3(*3*), GIANA(*4*),GLIPH(*5*)) and epitope annotation models (TCRex (*6*), ImmuneWatch™ DETECT, NetTCR(*7*), mixTCRpred(*8*)), however thus far these approaches have had limited adoption for single-cell data.(*5–8*)

The current state of the single cell field uses standardized pipelines and focuses on interrogating unsupervised-based annotations. However, this ‘GEx-centric’ approach may not be the most ideal methodology for every field. With T cells the critical interrogation challenge involves the analysis of single-cell data with not one but two key data levels. Conventionally, data analysis pipelines have been developed for bulk TCR-seq (*9*) and GEx experiments independently. Due to the historical development of GEx and TCR-specific analysis tools, there is a disconnect between these two data levels as they are analyzed separately rather than concurrently. Some efforts have been made to leverage both modalities, including CoNGA(*10*) and mvTCR(*11*). Those approaches that do integrate both levels build on the assumption that they are alternative representations of the same information, i.e. one TCR translates to one function(*10*), and does not consider the dynamic states of T cell sub-population(*12*). Most analysis pipelines simply extend on the existing methods for GEx-centric analysis (e.g., Seurat(*13*), scanpy(*14*), DALI(*15*)), with the TCR often seen as a secondary aspect to annotate the gene expression clusters by using is as an index marker for clonal expansion (*16*).

The GEx-centric approach therefore relies on the accuracy of T cell phenotyping across the unsupervised clustering. However, T cell sub-population differences are driven by subtle dynamic variations(*17*). This information may not be captured in low-dimensional projections of the GEx matrix, which is used to inform clustering of cell populations. (*16*)However, there has been little interrogation using the captured TCR to confirm the accuracy of the unsupervised T-cell subtype annotations.

Overall, the current GEx-centric analysis is currently ignoring a wealth of data within the captured TCR-seq layer. Thus, we aim to create a ‘field-centric’ methodology we defined as ‘TCR-first’ to overcome the generic GEx limitations. Here, we compiled a T cell atlas of 12 publicly available datasets containing ∼500,000 cells with both TCR and GEx and aimed to determine the utility of a TCR-first analysis. This required the development of a novel software tool coined ‘Single-cell T-cell receptor and Expression Grouped Ontologies’ (STEGO.R) to resolve many of the described analysis issues. Switching to a TCR-first approach, allows identification of (1) dynamic T cell states within a single clone across treatments, (2) epitope/disease-specific signatures, and (3) functional similarity between T-cells from different lineages. These findings resulted in a unique single T cell atlas, covering the general T cell phenotypes (Th1, Th2, CD8+ effector etc.), as well as state specific models e.g., cell cycling, immune checkpoint, cytotoxicity, and Th1 cytokine expression along with a TCR-seq based annotation. Overall, we demonstrate that it was beneficial to have ‘field-centric’ analysis consideration with this TCR-first approach.

## Results

### Current depth of TCR-seq analysis with GEx-centric approach

To showcase the current limitations of the GEx-centric approach in T cell research, we briefly summarise how the TCR information was described in each of the 12 papers included in the atlas (**Extended Table M1**). Briefly, 11 of 12 were analysed in R using Seurat with ten using scRepertoire that categorised the data into count categories of clonal expansion; none of the papers considered clonal expansion normalised as frequency. GSE185659(*18*), a transplantation focused dataset pre- and post-glucocorticoid treatment, was the only one to compare the top four clones compared to the remaining cells. Two studies GSE180268(*19*) and GSE145370(*20*) used the trajectory modelling of Monocle3(*21*). Only GSE180268 head and neck cancer T cell describing them transitioning from stem-like to intermediate profile and finally terminally differentiated state. Overall, the TCR-seq was underused across the 12 studies as the typical GEx-centric approach does not allow for in-depth interrogation of the TCR.

### T cell atlas identified background data was needed to improve FindMarker enrichment

The datasets were combined into a single atlas, where Harmony was used for the batch correction. We also limited the data to the TCR-GEx as these were high confidence T cells. Additionally, we hypothesized that combining the 12 datasets would improve overall annotations, especially with identifying rarer T cell phenotype and/or study-specific signatures. Originally all the datasets were analyzed separately; however, we identified a key issue regarding the process when utilizing the Seurat “FindMarker”, which was found to be dependent on similarity to the background.

To illustrate the annotation issue, we selected to use the lung transplantation (LTR) dataset (GSE185659(*18*)). This was one of the few non-cancer datasets, and therefore should have a distinct profile from the cancer-based disease settings. We aimed to identify if having a larger background dataset, would drive out disease specific signature in LTR. The top expanded clone TRAV8-3.TRAJ17 CAVGASKAAGNKLTF & TRBV6-5.TRBJ1-5 CASRRTGRNQPQHF was selected as it was present in acute cellular rejection (n=192) and post glucocorticoid therapy (n=2) for a pairwise analysis. This clone appeared to respond to therapy as it was depleted after glucocorticoid treatment. We compared this clone to the remainder of the LTR dataset (**Supplementary Figure S1A)** and the T-cell atlas (**Supplementary Figure S1B**). The clone profile vs the LTR dataset showed a more generic cytotoxic (GZMB) and activation profile (several HLA-DR transcripts). When this clone enrichment was done with the T cell atlas identified several district transcripts including Metallothionein 1E (MT1E), which appears to have a role as a regulator of T cell function in certain T cell subsets (*22*). MT1E was not detected to be different in the original analysis as it was widely expressed LTR dataset (**Supplementary Figure S1Aii**). To showcase that this was not because of a sequencing bias we visualized that MT1E was expressed to varying degrees across all 12 studies (**Supplementary Figure S1C**). This interrogation highlights the issue of similarity to background, which may mask important disease-specific signals. Therefore, we have made our T cell atlas available to reduce background issues that can occur in small and/or sorted datasets.

### TCR-first approach identified novel and dynamic signatures from a single clone previously hidden from the GEx-centric approach

To demonstrate the phenotypic heterogeneity of T cell clones with identical T cell receptor sequences, we used STEGO.R to assess the dynamics of the most prevalent clone in the 12 datasets (*18–20, 23–33*). The most hyperexpanded clones originated from the large granular lymphocyte (LGL) leukemia dataset, a T cell cancer (*29*). The original purpose of the LGL study was to block CD52 and cause apoptosis of the over-proliferated T cells. While some cases fully responded to treatment, the anti-CD52 (alemtuzumab) only caused a partial response in some patients including UPN4. For this analysis we focused on the partial-responder UPN4, with the aim to identify potential reasons for the lack of response.

UPN4’s highly expanded clone (n = 36,846), TRAV12-2.TRAJ9 CAATTGGFKTIF & TRBV20-1.TRBD2.TRBJ2-4 CSATEGNIQYF was likely driving their T cell malignancy. The clone featured a predominant CD8+ effector-like phenotype that contained a mixture of phenotypes based on our T cell functional annotation model (**Figure 1A**). Despite the prominent CD8+ phenotype, the standard unsupervised cluster-based annotation labeled several clones as CD4+ in pre: 5.7% and post: 2.6%. In contrast, our improved semi-supervised annotation model identified only pre: 0.43% and post: 0.07% with the CD4+ marker (**Supplementary Table S2**). The semi-supervised method thus had more accurate CD8+ annotations with ∼13x (pre: 5.7/0.43) and ∼37x (post 2.6/0.07) fewer incorrect CD4+ assignments (**Figure 1B**). To confirm the robustness of the semi-supervised annotation strategies, we analyzed the other top 50 clones with a threshold >95% for the CD8 marker. The semi-supervised approach identified 43 of the top 50 clones as CD8+, while using the unsupervised strategy only six clones met the 95% expression threshold. When annotating these CD8+ clones with the semi-supervised, they had fewer CD4+ miss-calls compared to the unsupervised approach (**Extended Results R1**).

**Figure 1.**
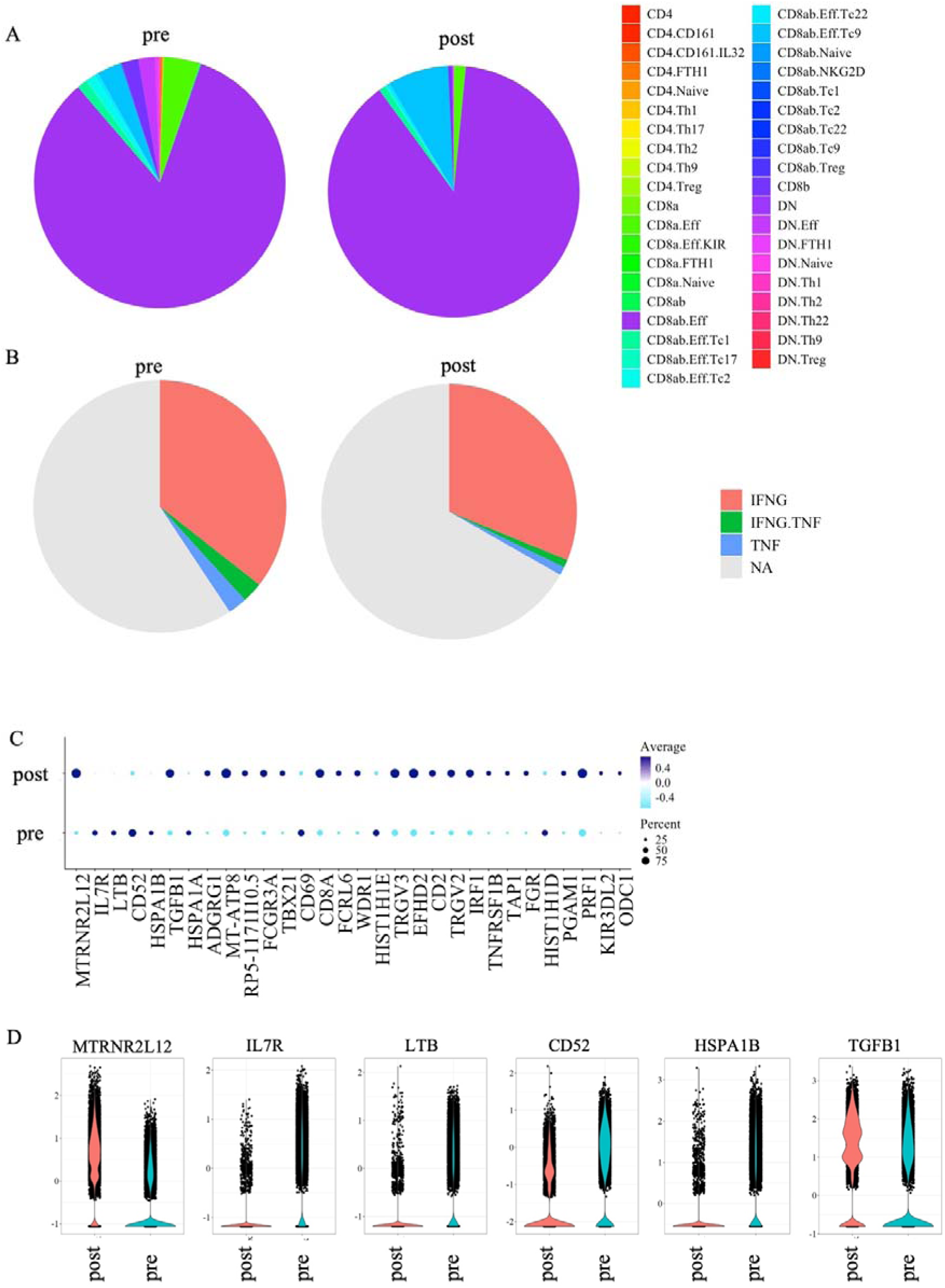
TCR-first approach identifies panel of genes post anti-CD52 from the malignant T cell TRAV12-2 & TRB20-1 clone from the partial responder UPN4. TRAV12-2.TRAJ9_CAATTGGFKTIF & TRBV20-1.TRBD2.TRBJ2-4_CSATEGNIQYF representing as (**A-B**) pie chart colored by the scGate modelling of (**A**) T cell functions using and (**B**) Th1_cytokines (IFNG and/or TNF). Comparing the pre- and post-anti-CD52 treatment showed as the (**C**) dot plot of the top 30 markers that were significantly different using the FindMarker statistic, and (**D**) the top seven genes expression represented as the violin plots of the post- and pre-treatment. For the violin plot, each dot represents a unique cell.

The TRAV12-2/TRBV20-1 clone CDR3 nucleotide sequence with 36,838/36,846 clones had the same nucleotide arrangement, TGTGCCGCGACTACTGGAGGCTTCAAAACTATCTTT and TGCAGTGCTACCGAGGGAAACATTCAGTACTTC, suggesting one originating event. To further characterize the TRAV12-2/TRBV20-1 clone through the measured gene expression of IFNG and TNF. The function of a T-cell is often assessed by the cytokine panel of Th1-based molecules IFNγ (IFNG) and TNF in experimental assays. Based on associated transcript markers, we identified that this T cell clone had a mixture of Th1 cytokine expression with subpopulations either being IFNG+TNF+ (3.8%), IFNG+ (46.3%) or TNF+ (2.6%) (**Figure 1B**). This illustrates how a single T cell clone is potentially differentially responding and depending on which cytokines they co-express and is reflective of what we observed in functional experiments. However, using known markers did not reveal substantive differences from either the population T cell phenotypes or Th1 cytokine expression pre- and post-anti-CD52 therapy that could explain the reason for the partial response.

To understand the impact of the anti-CD52 treatment on this T cell clone, we compared the transcriptional profile across time points (pseudo bulk analysis). The top transcript associated with post-treatment was MTRNR2L12 an emerging pseudogene that appears to have some relevant to CD8+ biology (*34*) (**Figure 1C; Supplementary Table S3**). There were also several transcripts with lower expression in post-treatment including IL7R, LTB, HSPA1B and CD52; the latter transcript’s protein was the intended target of the anti-CD52 treatment (**Figure 1D**). We can thus theorize that the reduced and lower expression of these genes may possibly explain the partial response of UPN4. This insight could only be found in the TCR-first approach as it was missed in the original GEx-centric analysis due to unsupervised global annotations and not interrogating specific clones.

### Identifying epitope-specific TCR and/or biomarkers that warrant functional validation

Some studies use tetramer sorting and various markers to identify clones and/or biomarkers of interest. The clone(s) of interest may not necessarily be the most expanded but are identified across different conditions with a distinct transcriptional profile. To illustrate this concept, we focused on a head and neck squamous cell carcinoma (HNSCC (*19*): GSE180268) dataset and used the remaining T cell atlas as the background to ensure signal enrichment when using the “FindMarker” statistic. In the original study, the authors state a clear narrative based on all single-cell and experimental data that PD-1 may be a potential target to boost the T cell immunity in HNSCC and concluded targeting PD-1 may be a good therapy to restore T cell function (*19*). The original analysis did not have an in-depth look at the TCR-seq to identify candidate TCRs that could have potential for screening, vaccine candidates and/or TCR-based therapies e.g., TCR-T.

Reanalysis of the T-cell receptor HNSCC data with STEGO.R revealed only a small number of public clones, which exhibited limited clonal expansion and lacked specificity towards disease (**Figure 2A; Supplementary Table S3**). Subsequently, we investigated the presence of clones shared across various treatments and conditions within individual patients and their clonal expansion. Specifically, as this dataset was tetramer sorted, we could deduce the TCR sequences likely able to recognize one of two HPV-epitopes HLA-A*01:01–HPV E2_329–337_ (peptide sequence KSAIVTLTY;KSA), and HLA-A*01:01–HPV E2_151–159_ (peptide sequence QVDYYGLYY; QVD)(*19*). There were clones that were KSA- and QVD-associated TCRs that were also identified in the PD-1 sort of primary tumor and metastatic lymph node (metLN) samples (**Figure 2A**). There was one KSA-specific clone in both HPV7 (**Figure 2B**) and HPV34 (**Figure 2C**). Additionally, HPV34 also had multiple QVD-specific clones (**Figure 2D; Supplemental Table S4**). Interestingly the QVD-specific HPV34 repertoire had many clones with TRAV21.TRAJ9 CAVDTGGFKTIF (n=11/27) and paired with different TCRβ arrangements.

**Figure 2.**
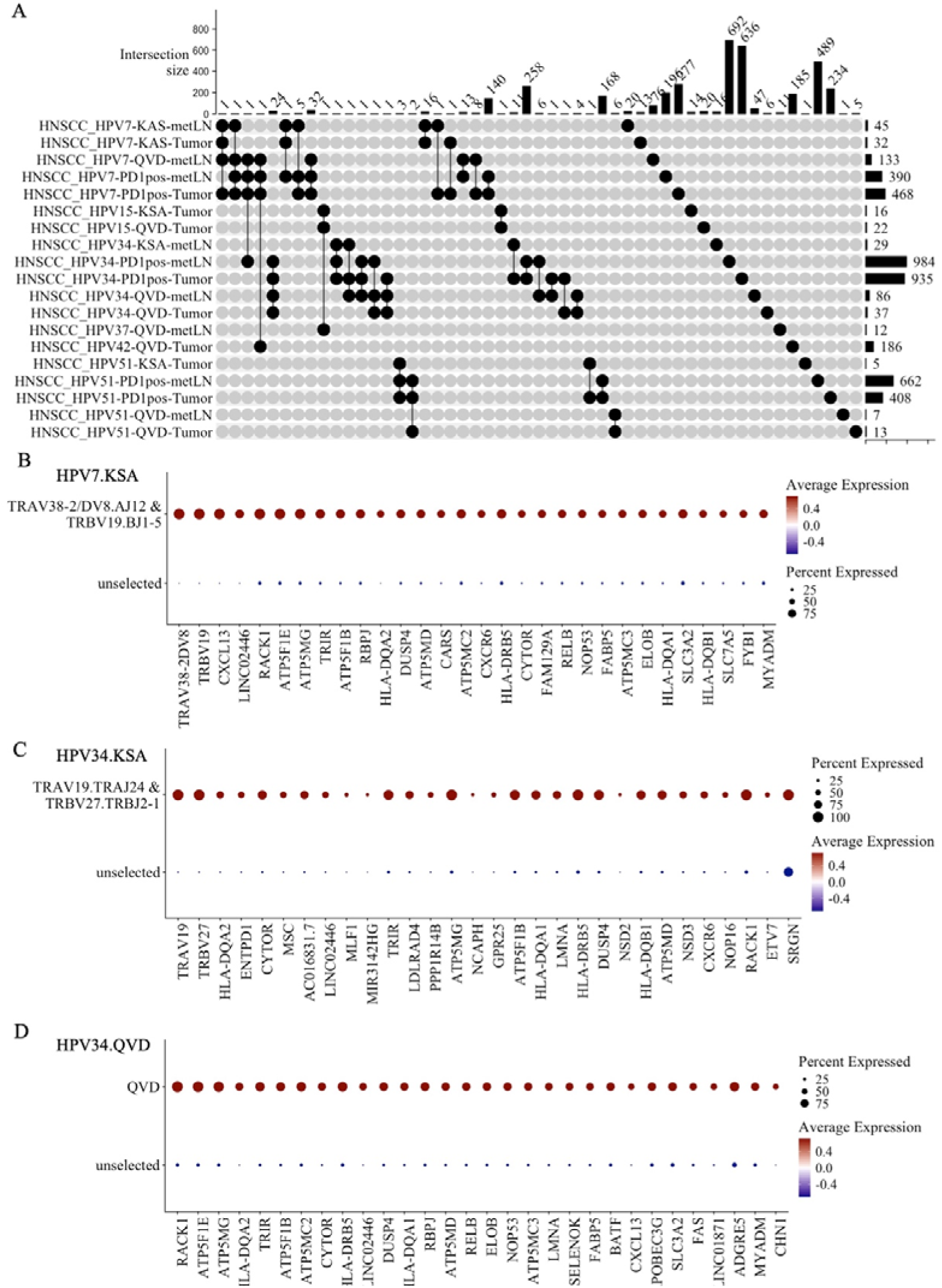
Re-examining the HNSCC identifies novel transcriptional signature of the HPV-epitope and PD-1 specific clones from HPV34. (**A**) Upset plot showcasing the unique overlapping sequences i.e., paired αβTCR with the CDR3 sequence, representing HPV7, HPV15, HPV34 and HPV51 with various sorting conditions (KSA, QVD, PD-1) and tissues (Tumor or metastatic lymph node[metLN]). The dot represents if a sequence was present in that sample. The lines connect the overlapping samples. The top bar-plot represents the number of overlap the samples. The right bar plot is the total unique sequences per sample. (**B-D**) Dot plots representing the top 30 significant genes associated from most significance (left to right) for (**B**) HPV7-KSA clones (TRAV38-2/DV8.TRAJ12 CAYNCPEPSDSSYKLIF & TRBV19.TRBJ1-5 CASSMLLNQPQHF) that overlap with both PD-1 from metLN and Tumor, (**C**) HPV34-KSA (TRAV19.TRAJ24 CALSGTDSWGKLQF & TRBV27.TRBJ2-1 CASSLSGTLGNEQFF) clone overlapping with PD-1 from metLN and Tumor and (**D**) pooled QVD-specific clones from HPV34 that were also identified in the PD-1+ from metLN and Tumor sorts.

We next compared if there were any significantly enriched genes common across the epitope-specific clones compared to the T-cell atlas for the above mentioned TCR’s of interest. There were 243 genes enrichment common to those TCRs (two KSA and pooled QVD clones), including CXCL13, LINC02446, RACK1, TRIR, RBPJ, HLA-DQA2 and DUSP4

(**Supplementary Table S5**). This indicates a T cell signature with consistent transcriptional markers from the epitope-specific T-cells that were also present in the PD-1 sort.

### Sequence similarity identifies both colitis- and melanoma-specific clusters

Despite the added benefit of utilizing the TCR to initiate the primary analysis, a common complication is that most TCRs are infrequent and private. In addition, all T cells express at least two distinct TCR chains (commonly αβTCR or γδTCR), which further complicates this analysis. To overcome these challenges, we propose the use of TCR clustering to group those sequences with high similarity into T-cell groups. Prior research has shown that high TCR sequence similarity within a population is often indicative of a common epitope response (*35*). Additionally, we also utilize the associated GEx and neighbor enrichment to further prioritize if the cluster is likely to have a unified function.

To demonstrate this approach, we reanalyzed a single-cell data set of colitis complication post-melanoma treatment as it was the only one with all four TCR chains (αβTCR and γδTCR) (*26*). This dataset comprised of eight colitis (C), six non-colitis (NC) patients and eight healthy controls (CT). For details on the overall TCR structure and common clusters, please see the **Extended Results R2**.

Interrogation of the alpha, beta, gamma and delta clusters revealed several intriguing patterns (**Supplementary Table S6 and S7**). One TRAV13-2-TRAJ45 cluster was present in all three conditions, with more sequences featuring in the CT and NC that had a mixture of effector CD8ab+ T cells as well as activation and cytotoxic features (**Figure 3A**). Ten of the 13 unique TRAV13-2-TRAJ45 sequences had significantly more sequence neighbors than expected (TCRdist >12.5 and p-adj <0.05; **Supplementary table S8**). These TCR paired with either TRBV4-1 or TRBV5-4. As these cells TCR pairing contained TRBV4-1 and TRAV13-2, it is possible that this cluster of TCR were CD1b-restricted (*36*).

**Figure 3.**
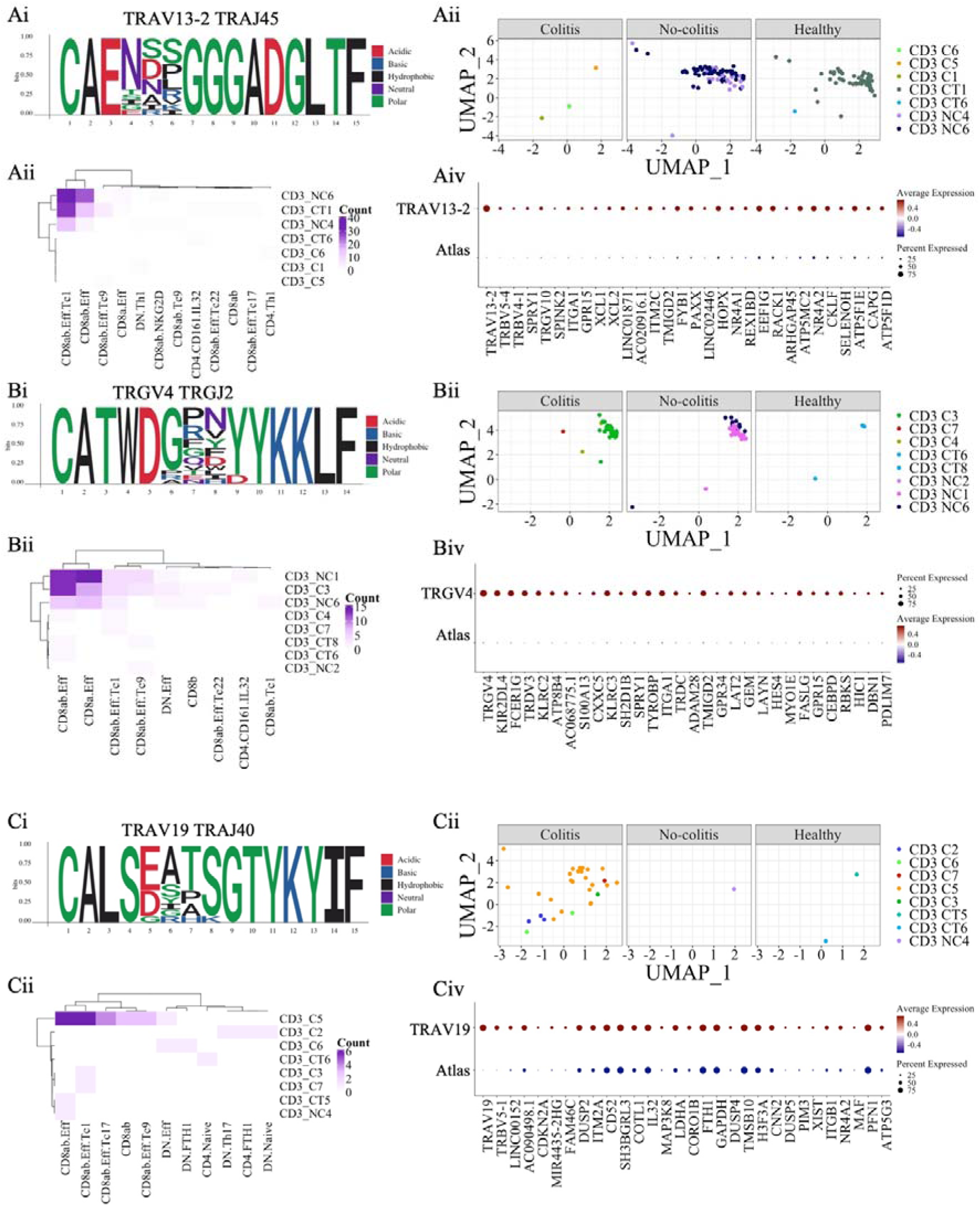
Clustering analysis of the colitis dataset highlighting disease specific clusters. (**i**) Motif of the cluster, (**ii**) Cluster on the UMAP plot coloured by individual and split by condition (normal controls, colitis and no-colitis), (**iii**) heatmap of the T cell phenotype vs the individual samples, and (**iv**) the top 30 transcripts enriched on this cluster compared to the remaining atlas (. C: colitis; NC: non-colitis; CT: normal controls. (**A**) Cluster 6 (TRAV13-2 TRAJ45) more common in NC and TC. (**B**) Cluster 8 (TRGV4 TRGJ2) more common in melanoma cases. (**C**) Cluster 9 (TRAV29/DV5 TRAJ40) more common in colitis cases. C, colitis; NC, non-colitis; CT, normal controls.

Interestingly, a TRGV4 gamma cluster associated with the C and NC cases, consisted of CD8ab+ γδ T cells with cytotoxic/activation status (**Figure 3B**). Again, we performed neighbor enrichment analysis, and found that eight out of 16 unique TRGV4 sequences in this cluster had significantly more sequence neighbors than expected (TCRdist >12.5 and p-adj <0.05; **Supplemental table S8**). As these TCRs had minimal presence in the healthy controls as well as being over-represented and expanded in the melanoma cases (C and NC) thus indicates that they are probably melanoma-associated TCRs.

Importantly, we also identify an alpha cluster with TRAV19-TRAJ40 that was over-represented in the colitis cases (**Figure 3C**). Within in this cluster, three TCR sequences had more neighbors than expected and were associated with the CD8+ effector phenotype (p-adj<0.05; **Supplementary Table S8**). This indicated that part of this TRAV19-TRAJ40 cluster may be colitis-specific.

The TCR beta chains contained fewer clusters of interest, which were mostly specific to a singular individual C3 (**Supplementary Figure S2**). The 11-mer TRBV6-2 (cluster 684) had four clones with significantly more neighbors than expected in the cluster, while the other 11-mer (cluster 377) and 12-mer (cluster 343) did not (**Supplementary Table S9**). Thus, only a few clones in one of the TRBV6-2 may have been colitis-specific or could have related to unknown infection.

Thus, our TCR-centered approach was able to identify disease-specific αβTCR clusters and γδTCR clusters that were not deducible from a GEx-centered methodology.

### Global TCR motifs associate with invariant T-cells and high generation probability

One of the benefits of combining the 12 datasets was to be able to identify common public patterns (*18–20, 23–33*). We describe some of the annotation differences observed in the dataset in **Extended Results R3** as well as the limited public clone interrogation in **Extended Results R4**. This below section focuses on public clusters common to all 12 datasets. There were three alpha clusters (**Supplementary Table S6**) and three beta clusters (**Supplementary Table S7**) common to all 12 datasets. The top public cluster had the classic TRAV1-2 with TRAJ33 that is the associated pattern for identifying mucosal invariant T (MAIT) cells (*37*). To identify the MAIT we compared the GEx to the TCR-seq (v_gene and j_gene). We found the TCR-layer was more accurately able to find the MAIT cells than the GEx alone (**Extended Results R5**).

The top alpha cluster had a junctional length of 12 amino acids and TRAV1-2 TRAJ33 gene arrangement and was present in 99 of 149 samples. This cluster had 1033 unique paired sequences with 1975 total clones. The DETECT tool predicted 67% (694/1033) of the paired unique sequences (Score >0.20) were likely to interact with MR1:5-OP-RU and indicated that the majority of this cluster were MAIT (**Figure 4A; Supplementary Table S10**). When examining the average GEx profile of this cluster not all cells expressed the commonly accepted MAIT markers, namely TRAV1-2 along with KLRB1 and SLC4A10 (**Figure 4B and 4C**). The two remaining alpha clusters, TRAV26-1 and TRAV3 were not functionally distinct from the background data. Additionally, as the significant genes associated with either were the V gene and generic T cell markers CD44 and CD28 (limited consistent enrichment), it suggests that there may be a diversity of functions for this cluster (**Supplementary Figure S3A and S3B)**. The IMW-DETECT did not identify any epitopes (>0.2) with present in either cluster.

**Figure 4.**
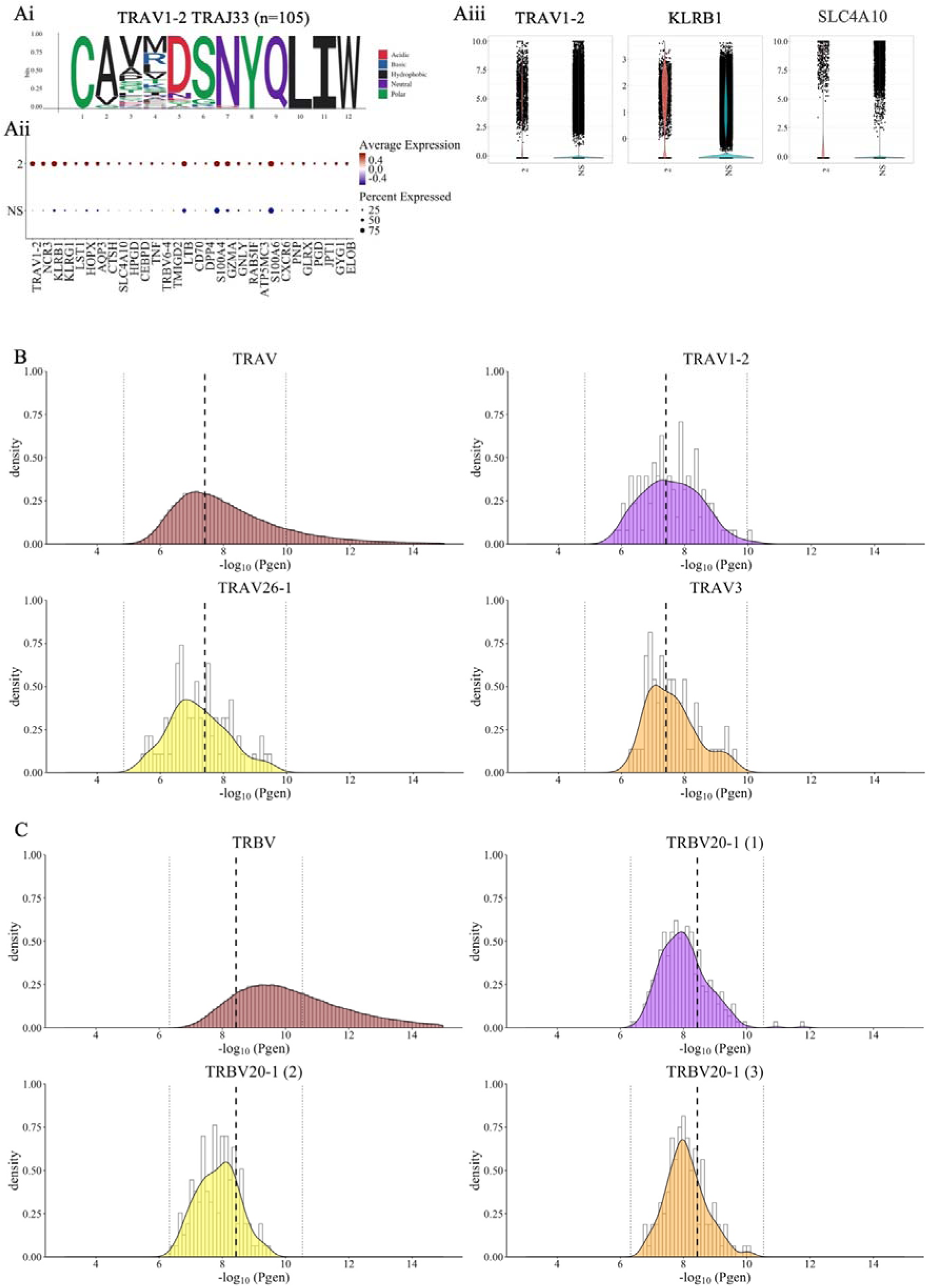
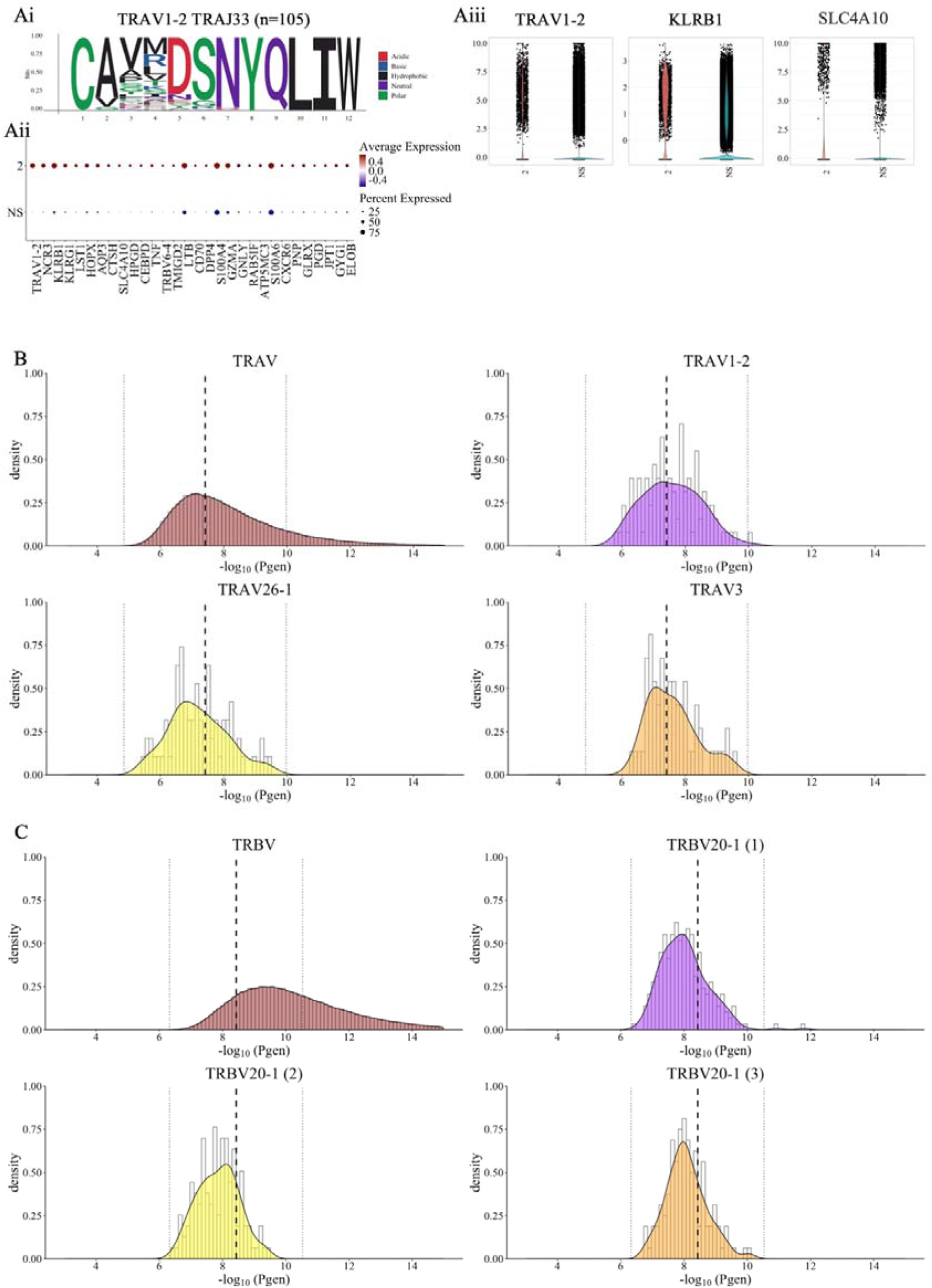
Global clustering analysis TRAV and TRBV clusters present in all 12 samples that had high generation probabilities. The top associated alpha cluster had the (**A**) TRAV1-2 TRAJ33 arrangement. This cluster was present as (**Ai**) a 12mer and (**Aii**) corresponding dot plot of the average relative expression that included the expected MAIT associated genes (Aiii) TRAV1-2, KLRB1 and SLC4A10. Addition we calculated the probability of generation with OLGA for (**B**) all TRAV probability of generation curve of the all sequences, TRAV1-2 TRAJ33, TRAV8-3 and TRAV27 and for (**C**) TRBV probability of generation curve of all sequences and the three TRBV20-1 clusters. (**B-C**) the middle dashed black line represents the geometric mean of the unique sequences, and the two dotted line represents one standard deviation from the geometric mean.

The three TRBV20-1 clusters had limited genes associated. All three beta-clusters contained TRBV20-1, had a junction length of 13 and lacked diversity gene inserts (**Supplementary Figure S3C to E**). FOS expression was noted in all three clusters, and this marker is associated with naïve cells (*38*). TRBV20-1 was the most common v_gene identified ∼17.1% of the total dataset (*39*). Of these TRBV20-1 sequences, the IMW-DETECT epitope prediction only three paired TCR were predicted to be MAIT cells (MR1:5-OP-RU; TRAV1-2 and TRBV20-1) and one TDLGQNLLY (Human mastadenovirus C; TRAV27 and TRBV20-1). This indicated that most cells had unknown epitope targets.

Next, we evaluated these six clusters recombination probabilities using OLGA(*39*). All clusters TCR arrangement were within one standard deviation of the geometric mean relative to the entire dataset (**Figure 4D and 4E**). Therefore, these sequences had a high generation probability, which may explain their prevalence across the different individuals and disease setting.

Lastly, we checked to see if any of the TCR sequence were more enriched in clusters than expected (TCRdist >12.5 and p-adj <0.05). The outcomes are summarized in **Table 1** and all data is present in (**Supplementary Table S11**). Interestingly, 82% of the TRAV1-2 TRAJ33 MAIT, 51% of the TRAV3 and 41% of TRAV26-1 unique sequences in the cluster had neighbor enrichment, indicating these clusters did not occur by random chance. On the other hand, few sequences in the TRBV20-1 clusters had more neighbor enrichment than expected (2.9%, 4.6% and 1.5%). Given the overall TRBV20-1 profiles with few associated differential transcripts and lack of neighbor enrichment, these T cells are probably the result of random chance.

**Table 1.** Summary table of neighbor enrichment for the six common clusters across the 12 studies.

| Cluster | Unique sequences in cluster. | Unique sequence found in enrichment and cluster. | Significant enrichment (p-adj <0.05) |
| --- | --- | --- | --- |
| **TRAV1-2 TRAJ33 (MAIT)** | 105 | 90 | 87 |
| **TRAV26-1** | 78 | 68 | 32 |
| **TRAV3** | 61 | 48 | 31 |
| **TRBV20-1 (1)** | 239 | 152 | 7 |
| **TRBV20-1 (2)** | 130 | 85 | 6 |
| **TRBV20-1 (3)** | 132 | 78 | 2 |

## Discussion

T cells are a complex cell type to interrogate due to their two-dimensional diversity in both the GEx and TCR. Under the current paradigm, GEx information is conventionally used to identify cell type clusters, which are subsequently labeled with features of the TCR. To shift the process to a TCR-first approach required the development of a novel tool STEGO.R to uncover actionable TCR-specific features. Here we re-interrogated and integrated 12 publicly available T cell datasets into a single cell-based T cell atlas and need for sufficient background data to ensure enrichment of signal. This atlas can be freely integrated into novel studies to aid in T cell annotation and improving enrichment of signals, especially for rarer population/clone, even when TCR data is unavailable (**Extended discussion**). Overall, the TCR-focused approach was able to identify patterns within the single cell data that were previously not identified from a GEx-centralized approach.

We first focused on validating the utility of the semi-supervised based annotations. As during thymus development mostly restrict to either CD4 and CD8 that determines future epitope recognition i.e., CD8+ T cells and class I MHC or CD4+ and class II MHC (*40*). The vast majority of the top 50 most expanded clones identified were likely CD8+ T cells. When examining the unsupervised clustering, many of the clones spanned multiple clusters and leading to lower confidence of the cell’s major CD8 vs CD4 expression markers. The semi-supervised method had greater accuracy especially for the CD8+ cells (>95% expression in 43/50 cells). Additionally, it appears that the unsupervised clustering often misses small perturbations that are important for the T-cell phenotype and necessitates the presence of other T cells of different phenotypes (**Extended discussion**). While no modeling is perfect, the semi-supervised annotation strategy had greater robustness over the standard unsupervised clustering for T cells. We recommend adopting this annotation strategy for future T cells studies.

Nevertheless, using global annotation strategies may be limiting understanding of specific T cell clones. We theorized that a lack of detectible difference was due to small GEx subtleties that drive the T-cell phenotype that is not captured in PCA used to create the UMAP plot for visualization. To better characterize the dynamic nature of T-cells, one can focus on a single TCR-defined clone across multiple time points. We showcased this by interrogating a hyperexpanded clone from the partial responder UPN4 from LGL(*29*) and identified a potential explanation as to the lack of response to the anti-CD52 therapy (**Extended discussion**). Additionally we demonstrated how the biased approach of GEx-centric method potentially missed key biomarkers in HNSCC re-analysis with receptor for activated C kinase 1 (RACK1) and CXCL13 more robustly indicating dysfunctional T cells (*41–43*) than the assumed PD-1 and/or CD39, as suggested by the original study (*19*) (**Extended discussion**). Therefore, to capture the overall profile, it was necessary interrogate a single TCR from two time points or pool TCR compared to T cell atlas as the background to identify a panel of markers associated with the TCR of interest.

Finally, we observe significant deficiencies in the definition of GEx-based functional annotation for T cells. A notable example for inadequate annotation protocols in T cell subtypes rely solely on the GEx information level, e.g., MAIT cells identified by TRAV1-2 and SLC4A10 or KLRB1 (*44, 45*). The commonly used unsupervised annotation strategy comparing clusters identified only one study to have possible MAIT cells. However, when interrogating the TCR-seq with the expected TRAV1-2 and TRAJ33/20/12 arrangement, we identified that MAIT cells transversed most studies and were diffusely identified across each cluster. We interrogated the top TRAV1-2 cluster with TRAJ33 arrangement present across all studies. Combining the GEx, IMW-DETECT tool and neighbor enrichment indicated that that this cluster did not occur by random change and implicated that most cells were likely MR1-restricted. In addition to this improved identification of MAIT populations, we also noted that γδTCR could have similar profiles to the adaptive CD8ab+ αβTCR and therefore, we cannot assume all cells with the CD8ab+ are only αβTCR (**Extended discussion**). There is also emerging evidence that CD8ab+ γδTCR have the potential for peptide restriction (*46, 47*), and they should not be ignored based on outdated understanding of γδ T cells being only innate-like (**Extended discussion**). Overall, the TCR-seq was needed to have higher confidence in finding both MAIT and γδTCR populations. We strongly suggest future single-cell T cell research to include both GEx and TCR-seq with all four T cell chains as this will further improving the accuracy in defining T cell subpopulations.

The full study TRB cluster interrogation identified common TRBV20-1 clusters with various J genes and was also the most prevalent TRBV gene identified. Our analysis identified that these sequences had a high generation probability, limited transcriptional enrichment with some Naïve T cell markers, and most of the cluster had the expected number of neighbors with <5% were significantly more enriched than expected. There were also no consistent IMW-DETECT epitopes identified. In the literature, TRBV20-1 TCRs are often among the most common TRBV gene arrangement in a multitude disease-settings including COVID-19 (*48*), non-small cell lung cancer (*49*), pancreatic cancer (*50*), rheumatoid arthritis (*51*), and range of cancers and tissue compartments (*52*). However, as the T-cells atlas shows, these are highly frequent TCRs across a multitude of conditions due to high generation probability. Thus when these TRBV20-1 are found present or even over-represented within a set of samples or a specific disease setting, this may solely be a proxy for a lack of response as they reflect the naive repertoire , or could be a statistical artefact based on the high abundance of TCRV20-1 TCRs and the intrinsic high TCR sample variability. For future studies caution is recommended before concluding if such common high generation probability sequences are indeed disease specific. We recommend applying the following strategy: clonal expansion and associated GEx, neighbor enrichment, screening epitope specificity from public databases (VDJdb (*53*)) or using annotation algorithms (TCRex(*6*) or IMW-DETECT).

We highlight through our interrogation that the generic GEx-centric approach is not necessarily the most appropriate methodology for T cell single-cell data, as correct understanding requires the dual modalities. This interrogation importantly showcased the improved robustness of semi-supervised based annotations that better matched the T cell biology. Collectively, our analyses illustrate that by shifting the interrogation to the TCR repertoire, one can uncover novel insights into defining T cell subtypes, as well as the identification of disease-associated TCR patterns. Exploring the clustering with TCR sequence neighbor enrichment and epitope prediction modelling could aid in identifying TCR of interest with functional understanding gained from the associated GEx. Moving forward we recommend a move towards a TCR-first methodology that is facilitated by our STEGO.R application. We believe this paradigm shift to a TCR repertoire focused approach will enhance our understanding of T cell biology and push forwards novel therapeutic opportunities.

## Methods

### Public datasets availability

We selected 22 publicly available scRNA-seq with scTCR-seq for benchmarking based on a literature search, of which only 12 of the 22 datasets could be processed (**Extended Table M1**) (*18–20, 23–33*). The main issues for not being able to process the remaining ten datasets were due to missing information in the public repositories (i.e., no available gene expression [n=2] (*54, 55*), no TCR-seq[n=2] (*56, 57*)), data format issue (e.g., summarized TCR data[n=1] (*58*), cannot separate cases in merged files [n=2] (*59, 60*), incompatible format[n=3] (*61–63*)). The 10x Genomics data was formatted as either raw filtered files (barcode, features, and matrix) or .h5 file. STEGO.R can currently process the raw files or .h5 standard formats. There is currently no process to extract a usable file from the cloupe and/or vloupe, and therefore were not analyzable with STEGO.R. The filtered outputs of the barcode, features, and matrix with the filtered_contig were the most accessible to processing in STEGO.R.

### STEGO.R Quality control process

The 12 datasets were processed in STEGO.R following the workflow outlined in https://stegor.readthedocs.io/en/latest/ and made use of the organized directory folder set up. Briefly, each of the 150 individual file formats for the QC processes were standardized in STEGO.R (**STEP 1**). This included formatting the matrix and meta data with the filtered TCR-seq, as well as downloading the ClusTCR2, TCRex and TCR_Explore files. As the current 10x Genomics kit and pipeline does not include all four chains, manual processing of the TCR-seq meta data was required for the colitis complication to melanoma treatment data set (*26*) to merge the alpha-beta (αβ) and gamma-delta (γδ) TCR-seq files. The TCRex and ClusTCR2 files were processed in **STEP 2**. Each of the 150 matrix files underwent individual quality assessment using the standard Seurat process and the processed TCR-seq was added to the meta-data, with LGL-healthy6 failing the QC process (**STEP 3a**; n=149). These files were then merged using the command-line function (**STEP 3b**). To ensure all cells could be annotated, the files were reduced to the 5006 transcripts and restricted to the TCR-seq that based on the original combining of the datasets during the merging process. The files were annotated using scGate for each 62,000 per loop (8 iterations) for the annotating. The annotations included the T cell functions (**Extended Table M2**) and major, immune checkpoint (IC), senescence, Th1 cytokines and cycling. In addition, TCR-seq information was used to define MAIT cells (TRAV1-2 J33/J20/12), possible CD1b/c restricted, γδ T cells and remaining αβ T cells (**STEP 3c**). **STEP 3d** was used to remove specific cells from the file based on the meta-data. Steps 1, 2b, 3a, 3b and 3c have command-line scripts available. More details on the inner workings of STEGO.R can be found in **Extended methods**.

#### IMW-DETECT

ImmuneWatch (IMW)-DETECT (Version 1.0) tool (2024; Available at: https://www.immunewatch.com/detect) is a user-friendly program designed to facilitate the prediction of epitope-TCR binding for paired TCR. It starts from a TCR file containing a list of paired TCR alpha and beta sequences, which must include information on the V and J gene and CDR3 amino acid sequence. IMW-DETECT annotates every TCR sequence for the most likely epitope target. More information on how to use this tool and interpret the results can be found at .

#### ClusTCR2

ClusTCR2 is an R alternative to the python package ClusTCR (*2*), specifically developed to be applied to single cell TCR sequencing data. The original version of ClusTCR applies a two-step process to cluster large sets of TCR sequences. The first step involves encoding TCRs into a numerical vector, based on the physicochemical properties of the amino acids. K-means clustering is applied to the amino acid vectors to create large families of pre-clusters. During the second step, a hashing function is used to identify all pairs of CDR3 amino acid sequences with ≤ 1 Hamming distance mismatch. From these CDR3 pairs, a graph is constructed , and communities are detected using the Markov clustering (MCL) algorithm, which will be the final clusters.

Since ClusTCR2 was developed with the focus on smaller, single-cell datasets, the second step is sufficient and achieves more accurate clustering results compared to the two-step method. Therefore, ClusTCR2 excludes the amino acid encoding and K-means clustering step. As the meta-data the alpha and gamma (_AG) chains can pair with their respective beta or delta chain (_BD). As we used the v_gene, a gamma chain will never be included in an alpha cluster. In the analysis section we also have separated the alpha (A), gamma (G), beta (B) and delta (D).

### Analysis process

These 12 distinct conditions with 149 unique files from 90 individuals contained 493,784 cells having both the GEx and TCR-seq. We aimed to determine if the analyzing TCR-first approach would identify novel patterns that are not discoverable from a GEx-centric approach.

The analysis process was from the perspective of a TCR-seq first approach, while the GEx-centric approach was documented in the original publications and limited to the scRepertoire clonal expansion onto the Uniform Manifold Approximation and Projection (UMAP) with very limited interrogation of the TCR-seq data.

The TCR-first approach was able to summarize the total clone count (public vs private), sequence similarity and where applicable predicted in TCRex. Each of these sections could identify the overall signature from the scGate annotations as well as the transcriptional signature expression (over-represented in TCR/cluster/population) compared to the remaining cells within the Seurat Object. The transcriptional signature was found with the ‘FindMarkers’ function in the Seurat Package and visualized as a dot plot (usually restricted to the top 30 transcripts, based on statistical significance). The default threshold used was a log fold change of 0.25 and p-value < 0.05.

To speed up the process, we used the automated function to extract the summary files and corresponding functions. We used either excel or R code to filer the files to identify which of the TCR, clusters or epitopes to specifically interrogate. Therefore, we did not have to manually create every file, and if needed updated the formatting in the STEGO.R applications for high quality, publication ready files.

Finally, we quantified sequence neighbor enrichment through comparison with a baseline model that estimates the expected neighbor distribution in the repertoire. In brief, TCR sequences are shuffled at the level of the junctions and a background data set is constructed from the shuffled sequences. Sequences are selected in such a way that the background approximates the V gene frequency, CDR3 length and generation probability distribution of the input repertoire. Here, we used a 50x (relative to the size of the input repertoire) background to accurately estimate expected neighbor counts. Sequence neighbor enrichment is calculated for each clone using the hypergeometric distribution. Bonferroni correction was applied to account for multiple hypothesis testing. The thresholds we used for a distance <12.5 and p-adj 0.05 to be deemed statistically significant. For interpretation, those with significant neighbor enrichment indicates that the clustering did not occur by random chance. While the null hypothesis is that >0.05 indicates that the cluster occurred by random change and has lower weight for biological importance.

## Supporting information

Extended Methods, results and discussion

Other table

## Code availability

STEGO.R code is available on the GitHub repository (https://github.com/KerryAM-R/STEGO.R).

## Funding

This work has been made possible by grant number 2022-249472 from the Chan Zuckerberg Initiative DAF, an advised fund of the Silicon Valley Community Foundation. In addition, this work was supported by the Research Foundation Flanders [FWO: 1S40321N to S.V. 1S71721N to N.d.V.].

## Conflict of interest

BO, KL and PM hold shares in ImmuneWatch BV, an immunoinformatics company, which developed IMW DETECT.

## Acknowledgements

KAM would like to thank: Dr. Nicole A Mifsud, Dr. Martin Davey, and Dr. Alexandra Sharland for providing access to their BD Rhapsody datasets to refine QC pipeline. The authors would also like to thank Romi Vandoren, Sofie Gielis, Fabio Affaticati, and Briek van den Born for testing the app and aiding in the development of the STEGO.R tutorial.

## Contribution

K.A.M., P.M., N.V., and S.V. were involved in the conceptualized the formation of the TCR-first analysis approach. K.A.M designed the software (STEGO.R) and writing the original draft. K.A.M. created the T cell atlas resource. K.A.M., B.O., N.V. and M.H., validated the data analysis steps. N.V. and S.V. aided with designing the formal analysis within the manuscript and STEGO.R. P.M., B.O. and K.L. supervised the project. All authors provided edited the manuscript and provided a critical review on its contents.

## Data availability

The publicly available data was sourced based on the GEO numbers (**Extended Table M1**). This final single cell T cell atlas .rds object is available in the Zenodo repository https://doi.org/10.5281/zenodo.10809382. In addition to the full 1.13M single cell T-cell atlas, we also include a down-sampled .rds consisting of up to 500 cells per T cell function annotation. This down sampled object can be used to help improve the overall annotation for new single T-cell data, and will be particularly useful if the dataset from an individual sample, or restricted subtypes.

---

**Supplementary Figure S1.**
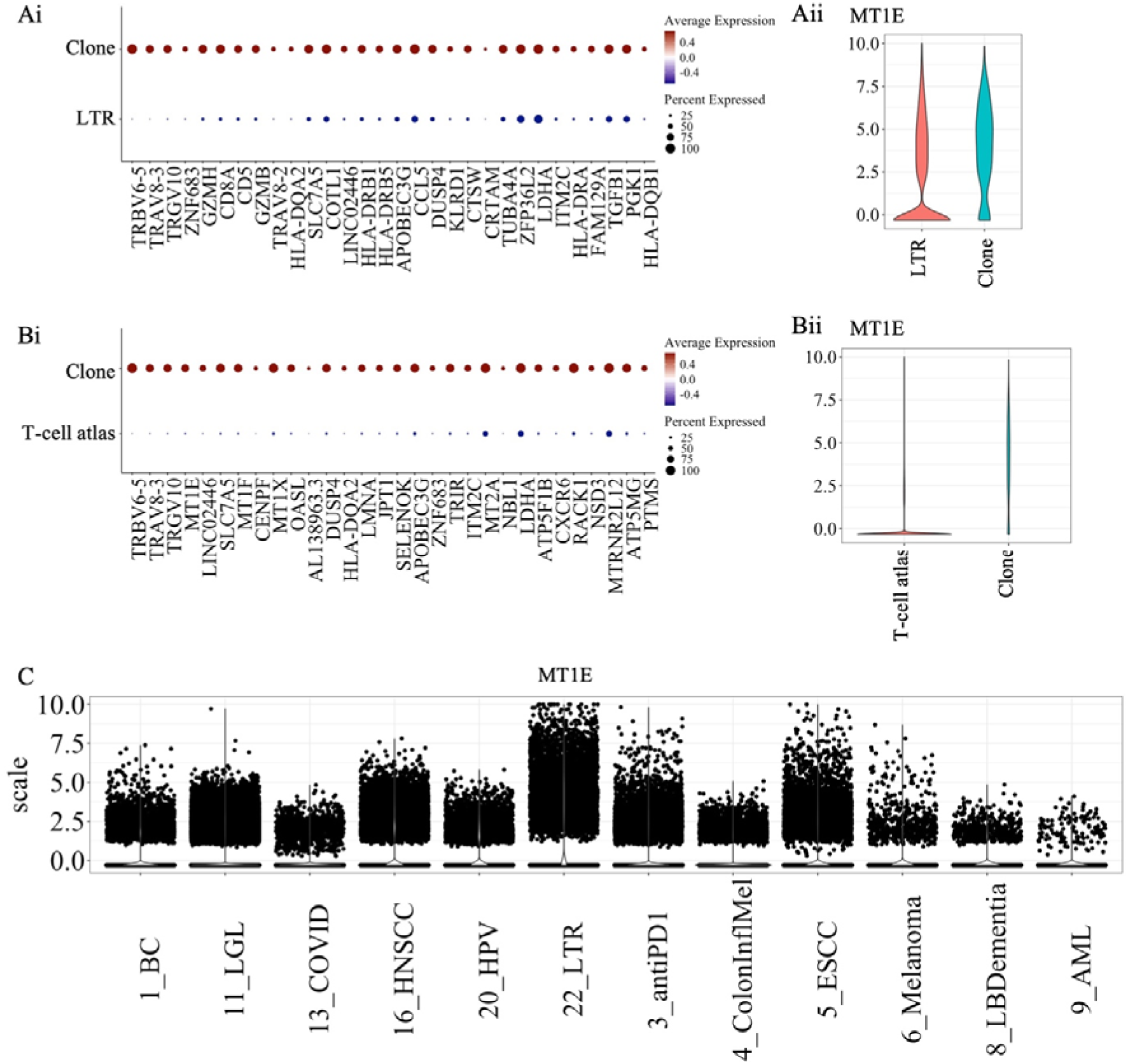
Genes enriched in the lung transplantation recipient (LTR) dataset. (**A-B**) Dot plot representing the normalized average expression and the approximant number of cells being expressed comparing the clone of interest TRAV8-3.TRAJ17 CAVGASKAAGNKLTF & TRBV6-5.TRBJ1-5 CASRRTGRNQPQHF to (**A**) the remaining LTR data and (**B**) remaining T cell atlas. (**i**) dot plots of the significantly enriched genes which showcases the percentage expressed and the relative expression. (**ii**) Violin plot of the MT1E transcript. (**C**) Expression of MT1E across the 12 studies that showcase the range of expression.

**Supplementary Figure 2.**
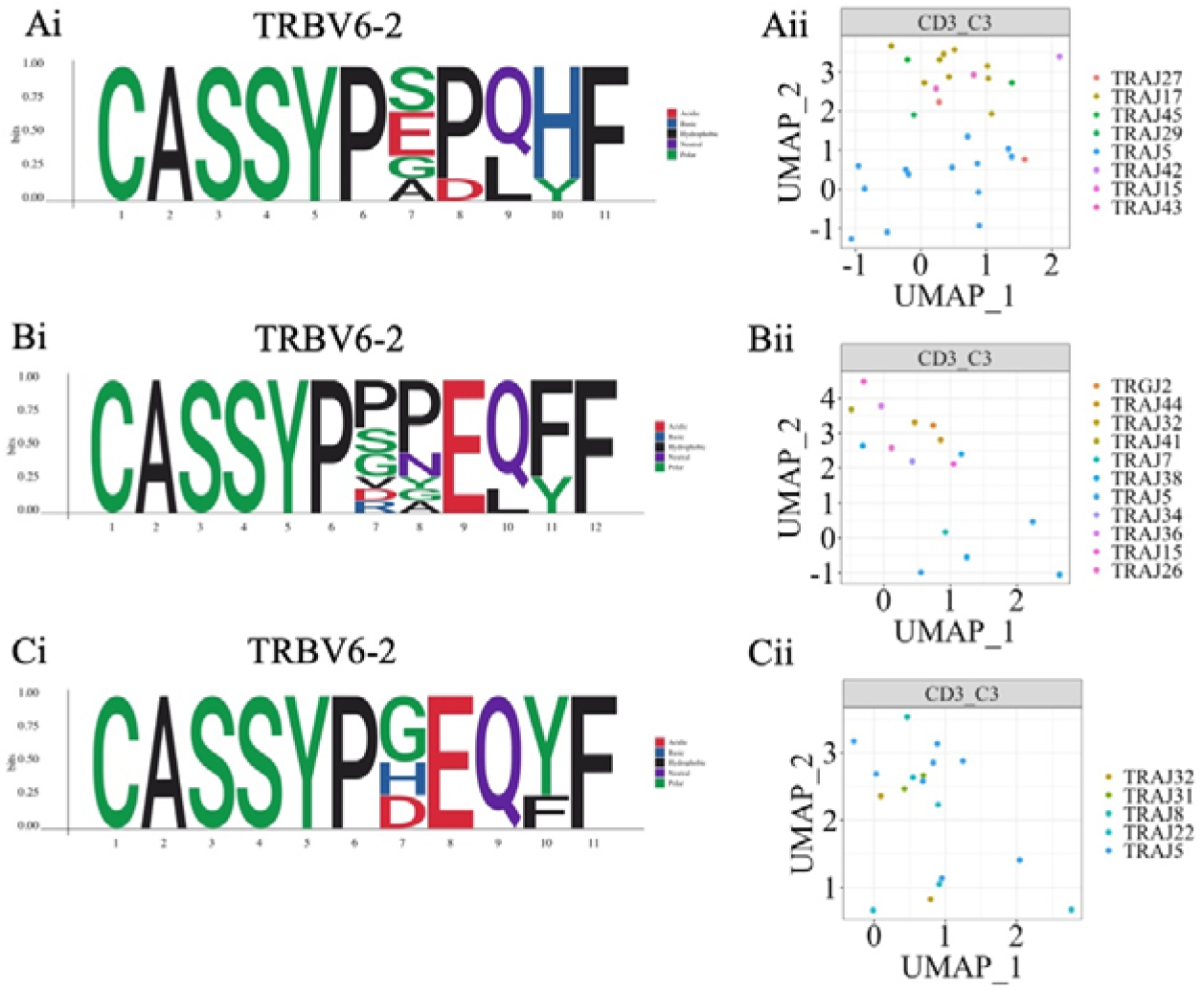
C3-colitis dataset showcasing three private clusters. (**A-C**) Three TRBV6-2 clusters from the C3-colitis individual. The three TRBV6-2 (**i**) motif plot are showcases as two (**A and C**) 11-mers and one (**B**) 12-mer. (**ii**) UMAP plot was split by the individual and colored by TRAJ gene. C = colitis.

**Supplementary Figure 3.**
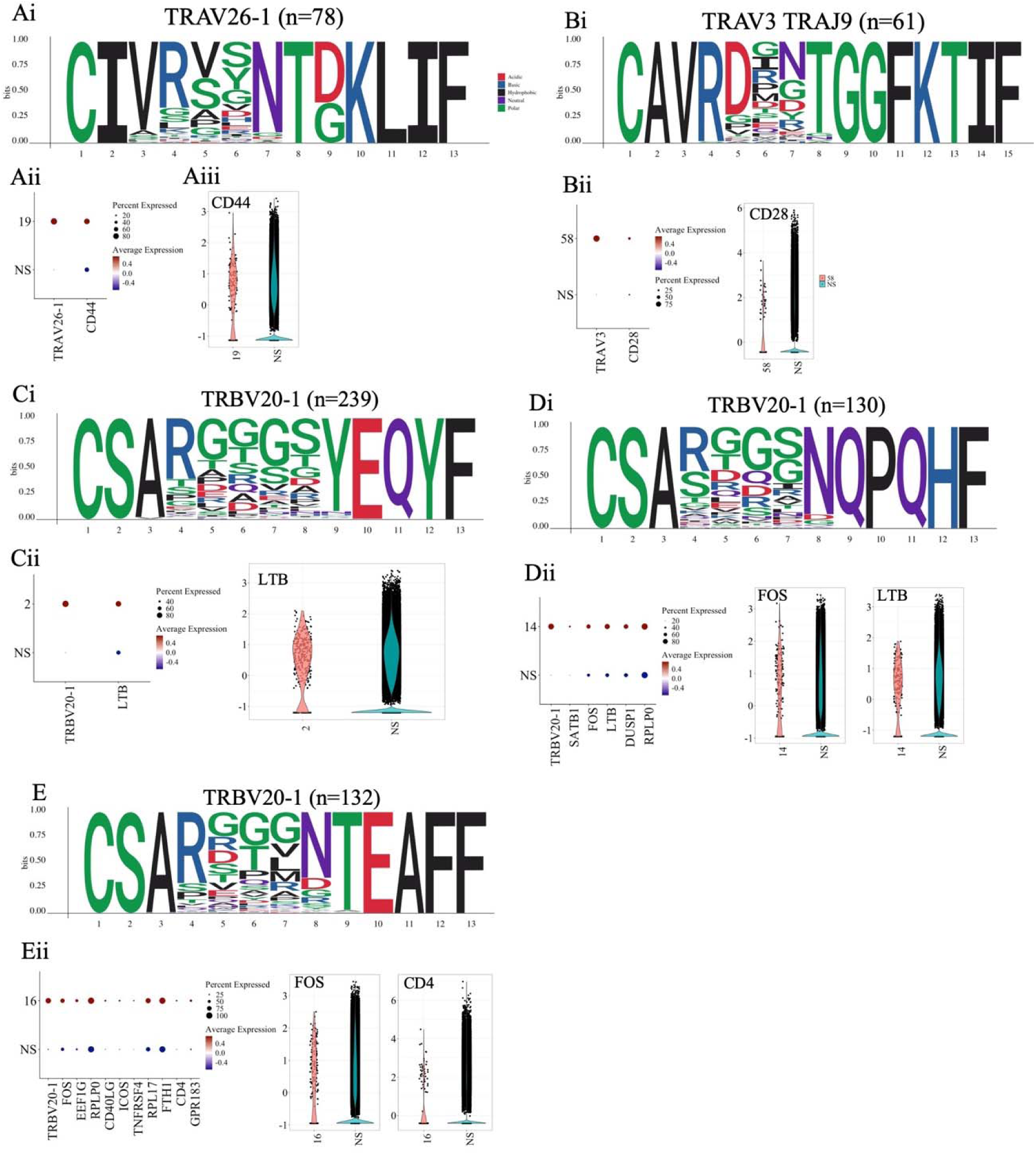
Global clustering analysis of two TRAV and three TRBV common clusters. Represent the remaining two (**A**) TRAV8-3 and (**B**) TRAV27 clusters as well as the (**C-E**) three TRBV20-1 clusters. The numbers in the brackets on the (i) Motif represent the total number of unique sequences; the number in the brackets represents the number of unique sequences present. (**ii**) dot plot and violine plot of some of the transcripts associated with the respective cluster.

## Notes

https://zenodo.org/doi/10.5281/zenodo.10809381

### Summary of Updates

Based on reviewer feedback from Nucleic acid research, we did a major revision of the current manuscript.

