## Extended Methods, results and discussion for "T cell receptor-centric perspective to multimodal single-cell data analysis"

#### Inner workings of STEGO.R

STEGO.R was created to aid in automate the T cell receptor repertoire analysis and prioritize TCRs in context of the gene expression from single cell experiments.

If a user has already processed the single cell data using R and scRepertoire proceed to step 3e, otherwise, start from step 1.

The program supports outputs of either Cell ranger (10x Genomics) or Seven Bridges (BD Rhapsody) (**Table M1**). As there are slightly different formatting between the technologies, they require distinct inputs.

Table M1. Input data required for 10x Genomics and BD Rhapsody.

| Source | Inputs |
| --- | --- |
| 10x Genomics | Features.csv.gz: Gene ID (SYMBOL)  Barcode.csv.gz: Cell ID (AGGATTT_1)  Matrix.mtx.gz: Gene expression.  Contig file (AIRR) format: contains the V(D)J sequences |
| BD Rhapsody | Features.csv.gz: Gene ID (SYMBOL)  Barcode.csv.gz: Cell ID (numeric)  Matrix.mtx.gz: expression captured  Sample_Tag.csv: multiplexing annotations  Unfiltered contig file (AIRR) format or Dominant (AIRR): contains the V(D)J sequences |

##### Install STEGO.R.

For details see the <https://stegor-documents.readthedocs.io>

##### Running the program.

require(STEGO.R)

runSTEGO() or STEGO.R::runSTEGO()

##### Setting up the project.

Download the Director_for_project that contains the directory structure of each project. Rename the “Director_for_project” folder as desired. For instance, “BC_2024” for the breast cancer dataset. The directory contains several folders that will be used for the processing (**Table M2**).

**Table M2. Structure of director and purpose of each folder**

| **File** | **Inputs** |
| --- | --- |
| 0_rawfiles | Copy the raw 10x Genomics/BD Rhapsody outputs. |
| 1_ClusTCR2 | Storage of the alpha-gamma (AG) or beta-delta (BD) clusters for each sample.  Store the merged AG_ or BD_ file for the clustering. |
| 1_SeuratObj | Contains the matrix file (cell x gene) and the meta data that contains the TCR information. |
| 1_TCR_Explore | Contains the (.csv) TCR_Explore files which can be used in the webapp <http://tcr-explore.erc.monash.edu> for visualizing the repertoire alone. |
| 1_TCRex | Contains the (.tsv) file that are in the required format for predicting epitope specificity in <https://tcrex.biodatamining.be> |
| 2_SCobj | Contains the Seurat objects (.rds) files. The user will save the individual QC files and the merged file and keep file (if required to remove sample issues). |
| 3_Analysis | This file stores all the outputs required for the STEGO.R analysis section.  This includes annotated_seurat.rds, AG_cluster.csv, BD_cluster.csv and TCRex.tsv  The folder also include .csv file for updating the ID’s as required. |
| custom_db | Contains nine folders that can be used by scGate for creating new annotation strategies. ‘master_table.tsv’ contains the ‘name’ and ‘signature’ required for all the annotation strategies. The specific strategies are in the files with the suffix ‘_scGate_model.tsv’. This suffix is required to identify each annotation label. Only use numbers and letters with no spaces for the file name. |
| Figures.Tables | Save the manual output files if desired |
| Prioritization | This directory will contain the automated outputs of the analysis for a first pass look over the results.  Contains the following sub-directories:  Clustering   - A - G - B - D   EpitopePred   - Can also be used for identifying TCR per annotation model   Multi   - PublicLike (for clones present in multiple samples) - Unique (clones present in one sample only)   ImmunoDom (single sample)  PolyClonal (single sample) |

##### Performing the QC process

STEP 0. Store all the raw file’s outputs in folder 0_rawfiles.

STEP 1. Upload the raw files into the QC section depending on the technology. Download each of tab outputs into their respective folders; 1_TCRex (TCRex.tsv), 1_SeuratQC (cell by gene matrix and meta.data file), 1_TCR_Explore (TCR_Explore.csv) and 1_ClusTCR (AG_clusTCR.csv and BD_clusTCR.csv).

Alternatively, this process can also be performed using the function in the **preprocessing.R** file. This process requires that the sub-folders (e.g., Sample_treatment) within the 0_rawfiles, need to contain the barcode, features, matrix and contig file.

STEP 2a. TCRex processing. Merge the TCRex files (if required), and upload (max 50,000 unique sequences) to <https://tcrex.biodatamining.be>. Download the output and save the **3_Analysis** folder.

STEP 2b. ClusTCR2 processing.

Merge the AG_ separately from the BD_ files. Load the merged file under the clustering inputs tab, click ‘run clustering’ and wait for ClusTCR2 to run. Download each of the AG_ and BD_ file to **3_Analysis** folder.

STEP 3a. Perform the Seurat QC for each file. Select the cut-off on sample mitochondria genes (mtDNA), ribosomal RNA (rRNA), feature counts. These cut-offs will change depending on the organism, and differences between focused panels (BD Rhapsody immune panel) and full transcriptome (both technologies). For instance, the default 10x Genomics parameters are <20% mtDNA, >5% rRNA, 200 to 2500 features. Low rRNA are likely sequencing issues. Higher mtDNA identifies cells that are damaged mitochondria and may represent a technical error, rather than disease specific features. Low features represent poor coverage, while higher feature counts likely represent doublets [doi:10.1186/s13059-016-0888-1]. before starting the process, the user adds in their file name, as it will be included in the ‘orig.ident’ column. The user will then select the number of principal components for dimensionality reduction (default: 15; usual range 10-15), as well as the number of distinct clusters. Once the QC of the Seurat object is finished, the meta-data containing the TCR sequences are added. The files is then saved in the 2_SCobj folder. This step is repeated for all files in your experiment. As there is a need to visualize the cut-off, this QC processing step was not automated. See <https://satijalab.org/seurat/> for details on the process.

STEP 3b Merging and batch correction. Merge the multiple Seurat files from the 2_SCobj. This process will restrict the number of features to a maximum of 5036 that was based on the 5000 most variable transcripts of the 12 datasets 5000 and the transcripts required for annotating. This step reduces the overall file size, enabling the ability to merge larger datasets, which may help decreasing computing time for the analysis. Save the ProjectID_merged.rds file in the **2_SCobj** folder. The merged file will then need to undergo batch correction with Harmony. This process follows the Seurat QC: find variable features, scale data, compute principal components, Harmony batch correction and UMAP dimensionality reduction. This batch correction is based on the ‘orig.ident’ column. Once completed, save the ProjectID_harmony.rds file in the 2_SCobj folder.

Step 3c. Annotating the Seurat object. Upload either the single sample file or merged file to annotate. Select the appropriate model, dependent on sequencing company and library size (e.g., all transcripts vs focused immune panel). These strategies have been modified for the BD Rhapsody Immune panel which has fewer genes and requires a high scGate threshold. Additionally, the current version also adds TCR-seq based annotations and needs to be added separately from the scGate annotation. We recommend the following annotation order: TCR-seq annotations, pre-defined scGate models and custom annotation. The final Seurat_annotated.rds file is saved to the **3_Analysis** folder.

After the refinement process, this should result in five scGate annotation models and a TCR-seq model for humans: T cell functions, major markers, cycling, senescence, immune checkpoint, as well as the TCR-seq based annotations for the invariant T cells e.g., MAIT, iNKT and distinguishing γδTCR and remaining αβTCR.

Custom annotation models:

1. Decide which genes to use for the annotation model.

2. Under the ‘marker check’ tab you can view if the genes are present via the feature plots and have been scaled. If not present, or lowly present relative to the CD8A expression, use with caution as they may miss many cells, which is the case with CD4 in many experiments.

3. In the master_table.tsv file add in the marker sets under ‘name’ and the gene ID’s under ‘signature’. The genes are separated by ‘;’ and each gene is considered as an OR variable in scGate (either can be present).

4. Create the requirements of each annotation (e.g., TcellFunction). The file includes the following headers ‘levels’, ‘use_as’, ‘name’ and ‘signature’. Levels column set as ‘level1, level2 … levelx’ acts like the gating strategy similar to flow cytometry and is considered as an ‘AND’ statement and if they are present which is designated as either positive or negative in the ‘use_as’ column. Update the suffix to the required annotation e.g., Early_scGate_model.tsv. the ‘_scGate_model.tsv’ is required to identify the files to add to the annotation model.

5. Update the name of the annotation model and run the model.

6. Check the locations of the annotations under the “UMAP check” tab.

7. Download the annotated file.

*Note: New annotations can be added to an already annotated file, as required.*

STEP 3d (optional). Remove unwanted aspects of the file. Even after the data has been process, the user may wish to further clean there file by removing samples based on any of the columns within the meta data table. For instance, in BD-Rhapsody datasets that use multiplexing, which can result in ambiguous origin with either multiplet or undetermined. These identifiers are located under the ‘Sample_Name’ column. This step can remove other unwanted samples as well (e.g., *NA* from the chain to restrict to both TCR and GEx). Save the file ProjectID_keep.rds to **3_Analysis** folder.

*Note: this step can be done before the annotation step.*

STEP 3e. converting scRepertoire files into the STEGO.R format. If users have already completed the QC process and have formatted the data in the scRepertoire formatting, this step will convert

##### STEP 4. Analysis

The analysis folder will contain several files required for interrogating the breadth of the repertoire including the annotated Seurat (.rds) file, clustering outputs (AG_ and BD_ .csv) and the TCRex (.tsv) outputs in the 3_Analysis folder. If required, for more complex experiments, the user can update the Update_ID.csv file, this file will match the ID used under the “Sample_Name” column. The “Sample_Name” column was added to the 10x Genomics pipeline, and is included in the “Sample_Tag.csv” file in the BD Rhapsody pipeline. The user can then change the ‘Selected Individual’, ‘Colour by:’ and ‘Split graph by:’ to the updated values as needed.

The user can upload the .rds file and it will automatically detect if the file was of BD Rhapsody or 10x Genomics origin, as well as the species. The BD Rhapsody detect if numeric were used in the Cell_Index, and the ending of ‘-1’ in the 10x Genomics. The species are detected from the scaled data for the presence of uppercase letters for the first three characters (*Homo sapiens* nomenclature) or uppercase first letter followed by lowercase letters (*Mus musculus* nomenclature).

The Analysis section has three main sections: overview, TCR and GEX, and prioritization.

##### Overview section

The overview section allows visualization of clonal expansion (TCR tab) and the gene expression (GEX tab). Unlike other programs/packaged (e.g., scRepertoire), which take the approach of performing the analysis of gene clusters annotations and then describe the TCR, we did not go into depth on this analysis approach.

The clonal expansion categories are based on the count category definitions described in scRepertoire (1): single (n=1), small (1<x≤5), medium (5<x≤20), large (20<x≤100) and Hyperexpanded (100<x≤500) and frequency categories: rare (<0.0001), small (0.0001<x≤0.001), medium (0.001<x≤0.01), large (0.01<x≤0.1) and Hyperexpanded (0.1<x≤1). This can be visualized on either a bar plot or overlayed on the UMAP. Additionally, the program introduces identifying the clonal overlap (downloadable Table) and visualized as an upset plot (<31 groups) .

The GEx tab allows the user to visualize the data on either the UMAP plot or as a proportion in a pie chart.

##### TCR → GEX

Unlike previous application and processes, the focus of STEGO was to interrogate the TCR repertoire and then identify their correlated gene expression. This section is separated into several sub-sections: Top clonotype, Expanded, ClusTCR2 and Epitope.

Common visualization and analysis for each section: summary table, UMAP of the selected TCR, pie chart of expression of the selected TCR, positive markers associated with the TCR relative the rest of the dataset, dot plot to display of the significant genes, and the over representation (all except Marker). The UMAP plots can be displayed as the overall sample or split by the group comparison “Include group comparison” to ‘yes’.

Over representation gene sets: There are 303 gene sets that are from various sources: Gene ontology biological pathways (GO BP), CellTypist, Reactome, KEGG, C8, MSigDB. Additionally, we also added in STEGO.R based gene sets (*see section 2.6.10 for details*).

Top clonotype: Summarizes the data based on the chain information with the default ‘vdj_gene_cdr3_AG_BD’ which ensures paired chain interrogations and can be changed as required. The default summary table is for the entire dataset, and if needed can be based on a single sample (set by: Display one individual? Yes, and then “Display one individual” to the selected individual). Once these variables have been decided, proceed to the stats tab, followed by the dot plot (can be restricted to number of genes), and the over representation analysis.

Expanded: This defines the expanded (Ex) vs the non-expanded (NEx) T cell clonotypes regardless of their specific sequences. This section can determine if the Ex have distinct signatures across samples, relative to the NEx of one sample. Alternatively, the user can also compare sample specific differences if they input the _Ex and _NEx into the ‘Samp 1’ and ‘Samp 2’ column. The default column to summarize is the ‘vdj_gene_CDR3_AG_BD’. The user will select the column to include “Sample column name” and select those to include in the ‘ID's to include’ box. In the side bar panel, the user can set Ex threshold based on frequency of the repertoire from in the “Cut off greater than’ column. Alternatively, they can select the minimum # of clonotypes in the “Cut off greater than”. The UMAP will display the Ex and NEx clones. Once these variables have been decided, proceed to the stats tab, followed by the dot plot (can be restricted to number of genes), and the over representation analysis.

ClusTCR2: This section focuses on the sequence similarity based on the ClusTCR2 package, which is the R based version of ClusTCR (2). The user will upload both AG_ and BD_ files separately. The side bar will contain two variables that the user to alter which cluster is displayed: “Clusters to display” (numeric value) and “Chain to display” (AG or BD). The order of the clusters is based on total number of nodes per cluster. The cluster motif can be visualized under the ‘motif’ tab which also displays what the Variable and junction genes association with the sequence and from what samples. Once these variables have been decided, proceed to the stats tab, followed by the dot plot (can be restricted to number of genes), and the over representation analysis.

Epitope: Epitope prediction from TCRex. There are up to 100 epitope models within TCRex based on single Beta chain, TRBV and TRBJ. The TCRex file adds in the epitope and pathology. The user can view the summary file based on three variables: function (Colour Pie by (hm = y-axis)), group (Split Pie by (hm = x-axis)) and individual (Selected Individual). The user can then impute the epitope information, which is in order of highest # of total clones. Once these variables have been decided, proceed to the stats tab, followed by the dot plot (can be restricted to number of genes), and the over representation analysis.

Marker: Identify TCR’s associated with a single or dual markers. This section uses the scaled data for the visualization and analysis. The single marker section was used to identify which TCR’s are associated with markers of interest. Set the threshold of ‘Marker +ve cut-off (>)’ by interrogating the violin plot that is separated by “Sample_Name” (**Figure M1A**). If the violin plot looks like **Figure M1B**, it is likely to be a technical issue rather than a true result. We noted that many of the cytokines had this pattern and were unlikely to be real, and therefore were removed from the annotation models. Once these variables have been decided, proceed to the stats tab, followed by the dot plot that can be restricted to number of genes, and the over representation analysis. The dual marker section follows the same process as the single marker section except two thresholds and selecting the quadrant of interest.

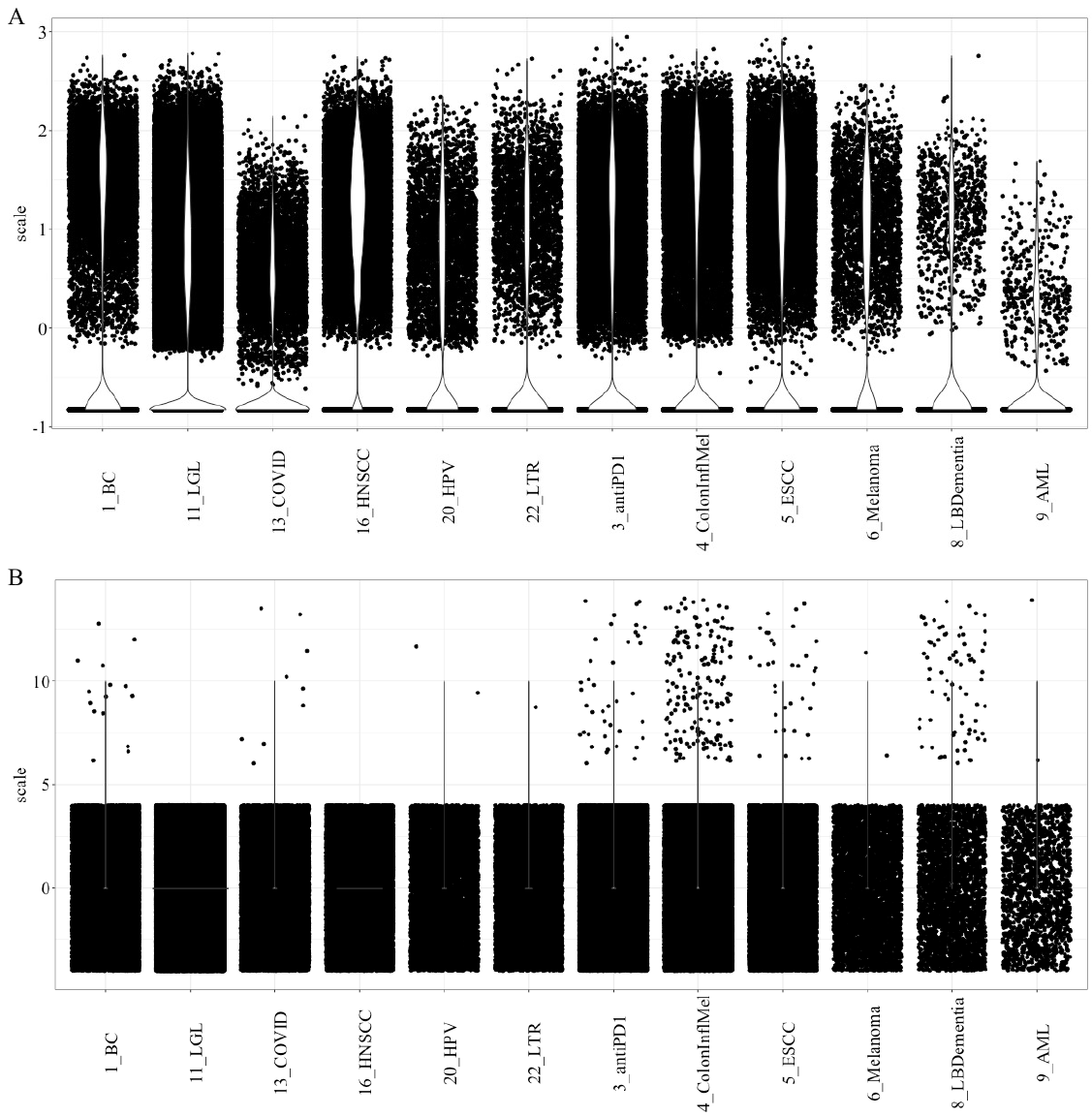

Figure M1. Violine plots showcasing quality and poor transcript expression. (A) high quality expression of CD8A, while (B) technical error of IL4. The x of the violine plot represent the unique studies and the y-axis is the scaling of the data.

##### Automating and TCR prioritization.

The automated strategy is found under the ‘prioritization’ tab. This section was split into three sub-sections: clonotype, cluster and epitope/annotation. The automated approach will use a minimum of three (>2) cut-off due to statistical requirement of the minimum sample size of three.

Clonotype

This section detects the type of data of single sample or multi sample based on the ‘Selected Individual’ variable (default: Sample_Name). The single cell dataset the user determines the percentage of the repertoire (side bar panel). The user can also increase the percentage until no ‘immunodominant’ clones are detected. The program will then compare the expanded vs the non-expanded clones.

When the program detects multiple samples, there is priority value to the clonotypes based on:

1/(total clonal frequency x total number of samples)

Therefore, the order of analysis is based on smaller priority to largest. The data frame is then subset to default to a minimum of two or more samples for the multi-sample analysis. The user can then set the threshold for the multi-sample priority where the summary is displayed in the table. This multi-sample detection section also identifies unique clonotypes (private), and the user can set the threshold (default >2).

The program will download the files in the ‘multi → Publiclike’ or ‘Multi → private’ folders respectively in the ‘directory_for_project’. The downloaded files include clone summary table and for each clone the find marker statistic table, dot-plot and over-represent table. The multi-sample will also download an upset plot if <31 samples are detected. The program will not download the overrepresentation table if no genes were found.

Cluster

The clustering section will require the user to upload both the AG_ and BD_ clustering file. The program updates based on the calculated priority score. This score is calculated the order based on:

1/(total clonal frequency x number of nodes x total number of samples)

Therefore, clusters with clonal expansion, more connections and present in multiple samples have higher weight. The user can change the threshold for both the separate AG and BD files. The user can update the threshold to decrease the total number of clusters analyzed.

The program will download the files in the ‘prioritization → clustering’ folder in the ‘director_for_project’ with output a clustering summary table and per clone will download the motif plot, find marker statistic table, dot-plot and over-represent table.

Epitope/Annotation

The user will upload the TCRex.tsv file that contains the beta and epitopes of interest. We recommend running both the ‘pathology’ or ‘beta’ (beta CDR3 sequence) for each epitope.

The function can also be used more broadly to the annotations that prioritizes the data based on a ‘group’ and ‘function’ e.g., Sample_Name and TcellFunctions.

The program will download the files in the ‘Prioritization → EpitopePred’ folder in the ‘director_for_project’ with output the epitope summary table, heatmap figure and per epitope/annotation will download a UMAP plot, find marker statistic table, dot-plot and over-represent table.

##### GEX → TCR

This section focuses TCR’s associated with certain annotations or specific markers (single or dual). As this was the previous approach to analyzing TCR per cluster annotation, this was not the focus on STEGO.R development. The automated extraction can be done under the Epitope/Annotation. For our purposes we used this section to validate which transcripts to use for the annotation modelling.

##### Development of STEGO.R transcript ontologies.

Given the lack of consistency in marker selection and limited pathway or ontologies based on the transcriptome, we sort to create transcriptional specific ontologies from the 12 publicly available datasets. The following markers were interrogated: SELL (Naïve-like), KLRG1 (Activation-like without TIGIT), FOXP3 (Tregs), RORC (Th17-like), CCR10 (Th22-like), IL21 (Tfh), IL4 (Th2), PDCD1 (PD-1 regulation of CD8+ T cell function), GZMK (memory-like), KLRB1, CXCR3, IFNγ (Activated), and KIR2DL3 (HLA-E presentation).

The significant genes associated with each marker were stored in “**Designing_STEGO.R_genesets.xlsx**” for each of the 12 datasets. The gene lists were imported into R and merged together. To create the associated transcripts, they must have been present in at least 50% of the datasets. IL4 was only present in one of the 12 datasets GSE161192, and therefore could not be created into a STEGO specific ontology. The following were selected as the gene cut-off(samples/total; number of associated genes): SELL (8/12; n=22), KLRG1 (8/12; n=10), FOXP3 (7/11; n=62), RORC (7/10; n=16), CCR10 (3/6; n=18), IL21 (7/11; n=14), PDCD1 (7/10; n=17), GZMK (7/11; n=26), KLRB1(6/12; n=6), CXCR3 (7/12; n=20), IFNG (7/11; n=30), and KIR2DL3 (5/8; n=13). This process can be replicated with the R script ‘GENESETS_Overlapping_Marker_lists.R’ and **Designing_STEGO.R_genesets.xlsx**” in the int/Global folder.

In addition to the above gene sets, we also added in NK-receptor based (common, inhibitory, activating, KIR-genes; 10.3389/fimmu.2019.01179) as well as granzyme genes and CD8+ T cell cytotoxic based.

### Extended Results

##### R1 Improve annotation strategy with semi-supervised approach.

The common annotation strategy uses the unsupervised annotation approach and requires determining the appropriate number of clusters. This can change with the addition of new data. With this combined dataset we set the resolution to 1 and resulted in 23 clusters. The find marker was used to identify the markers enriched in each cluster relative to everything else. If statements were then used to identify if the CD4 or CD8 marker were enriched for each of the 25 clusters. The feature expression of CD8A and CD4 shows various coverage for each of these major T cell markers within each cluster (**Extended Figure R1**). Therefore, neither CD8A nor CD4 were major contributing feature in the PCs that are used as inputs for the to create the unsupervised clustering.

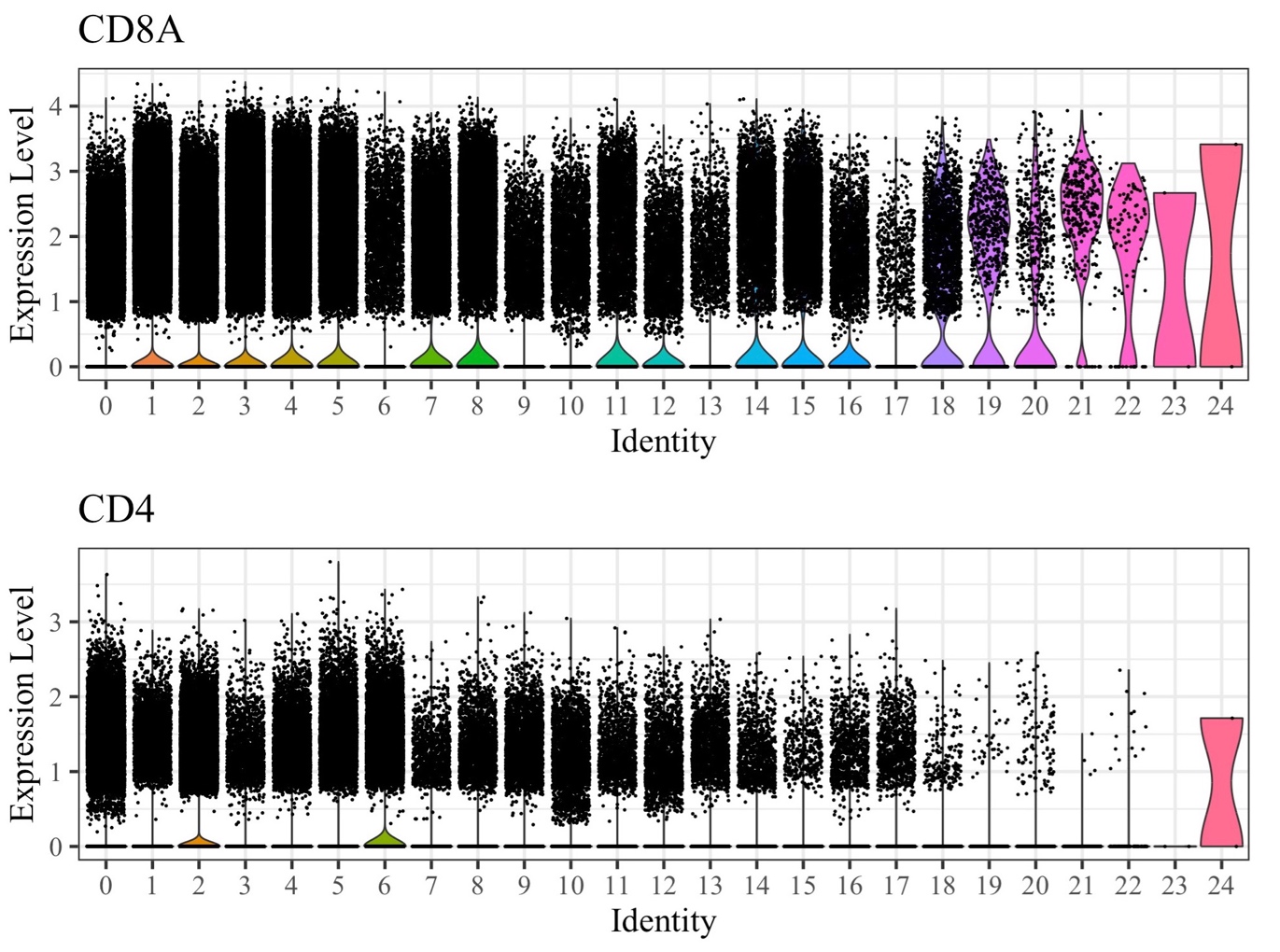

**Extended Figure R1. Expression of CD8A and CD4 per cluster.** The violin plots are split by the 25 clusters (0 to 24) and highlighting the (**top**) CD8A marker and (**bottom**) CD4 from the T cell Atlas ~500k cells. Each dot represents a unique cell. Each plot has the scaled expression from 0 to 5.

To confirm if the semi-supervised annotation strategy was more accurate than the unsupervised clustering, we interrogated the top 50 most expanded clones. For the CD8+ cells, the cluster-based approach identified a mean expression of 75% per clone (six clones with >95% of the total cells/clone expressing CD8) and the semi-supervised a mean expression of 94% (43 clones with >95% of the total cells/clone expressing CD8 (**Extended Figure R2; Extended Table R1**). Only one of the top 50 clones appeared to be a CD4+ T cell. This cell only had 50% coverage CD4 annotation in the semi-supervised approach, which it would have been mislabeled in the unsupervised with <1% deemed to be CD4 (**Extended Figure R3**). The lower confidence of the annotations is observed because the clones can span many of the unsupervised clusters and diluting the signal for assigning even the major T cell markers of CD4 and CD8 that determine function i.e., HLA/MHC specificity of class I or class I peptide presentation. This data identified that CD4 and CD8 were not among the major features to create the UMAP and had more diffuse expression across the clusters, indicating the issue in using the strategy for cell annotation. Overall, validating with the TCR-seq we were able to confirm that the semi-supervised annotations had higher confidence at calling CD8+ T cell assignment then the unsupervised strategy.

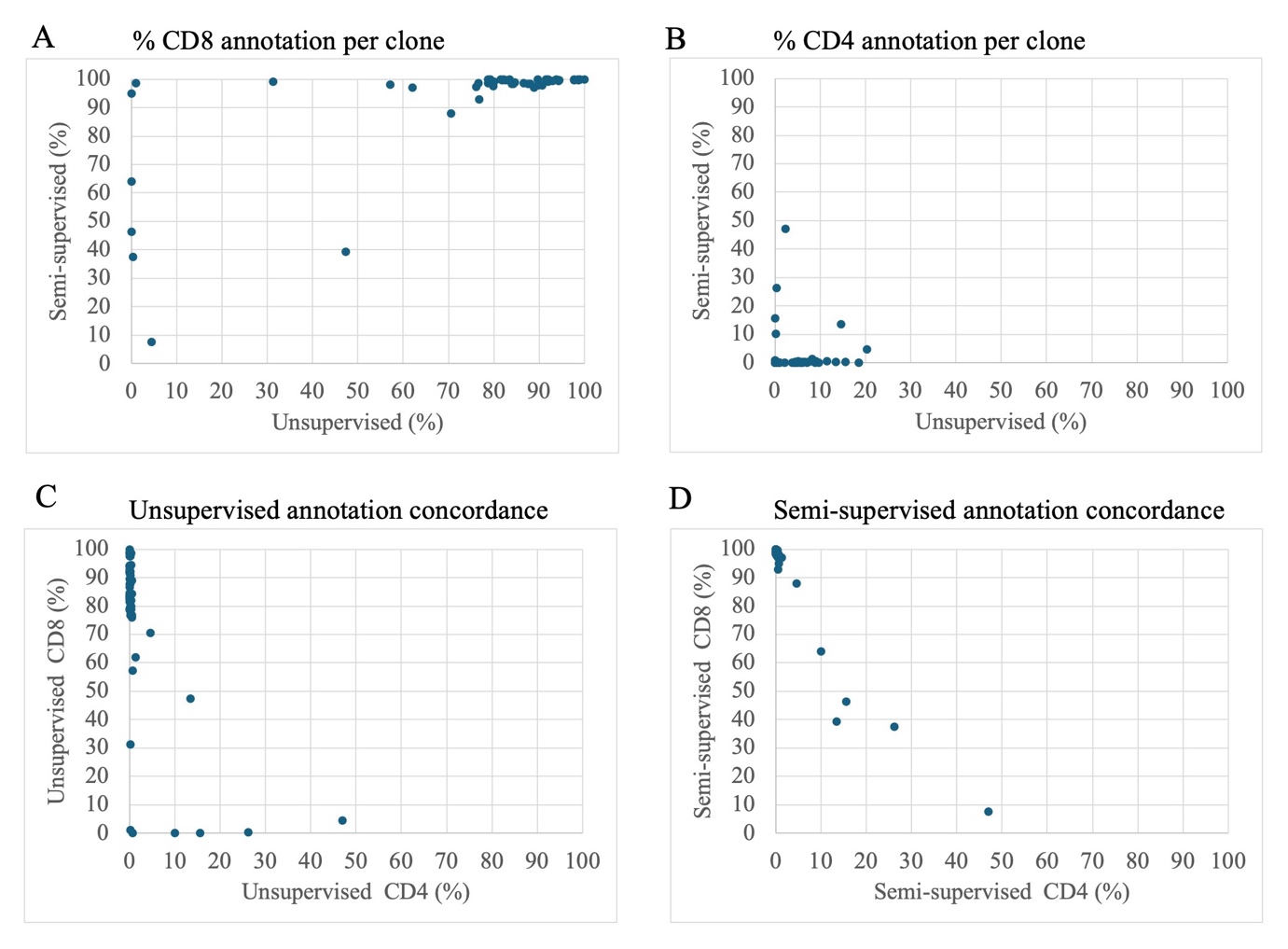

Extended Figure R2. Top 50 clones’ annotation of the major T cell marker. (A) CD8 and (B) CD4. We also compared the CD8 and CD4 percentages for (C) unsupervised and (D) semi-supervised clustering. (A-B) The x-axis is the percentage of cells expressing for the cluster-based annotation. The y-axis represents the percentage of cells expressing the CD8 semi-supervised based annotation. (C-D) x-axis represents the % CD4 expression and y-axis is the % of CD8 expression. Each dot represents a unique clone.

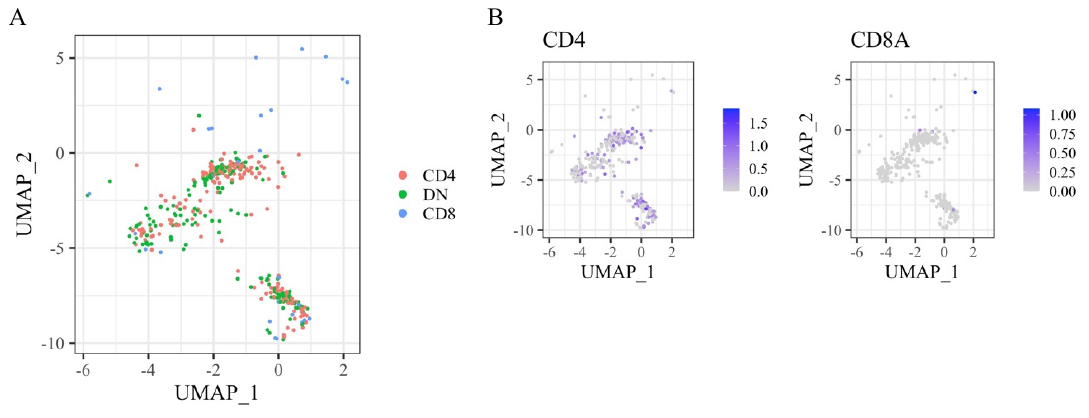

**Extended Figure R3. Likely CD4+ expanded clone: TRAV13-1.TRAJ28 CAATDSGAGSYQLTF & TRBV20-1.TRBJ2-1 CSAPLGTSNEQFF.** (**A**) The unsupervised clustered for this clone was identified in 11 different clusters. (**B**) Expression from left to right CD4 and CD8A.

##### R2 Identified limited public clones and common clusters in the colon dataset

From the colon derived T cells, there were 38,394 unique clones across the 22 individuals in the dataset with most clones being private. Indeed, only 49 clones were shared among two or more individuals (**Extended Figure R4**). Of these, 14 clonotypes were present in more than one disease state (**Extended Table R2**). Nine overlapped between the melanoma patients. Four overlapped in the normal controls and colitis, three of which were expanded in the colitis cases. Although the degree of expansion in these overlapping clones was minimal (n<10 per clone).

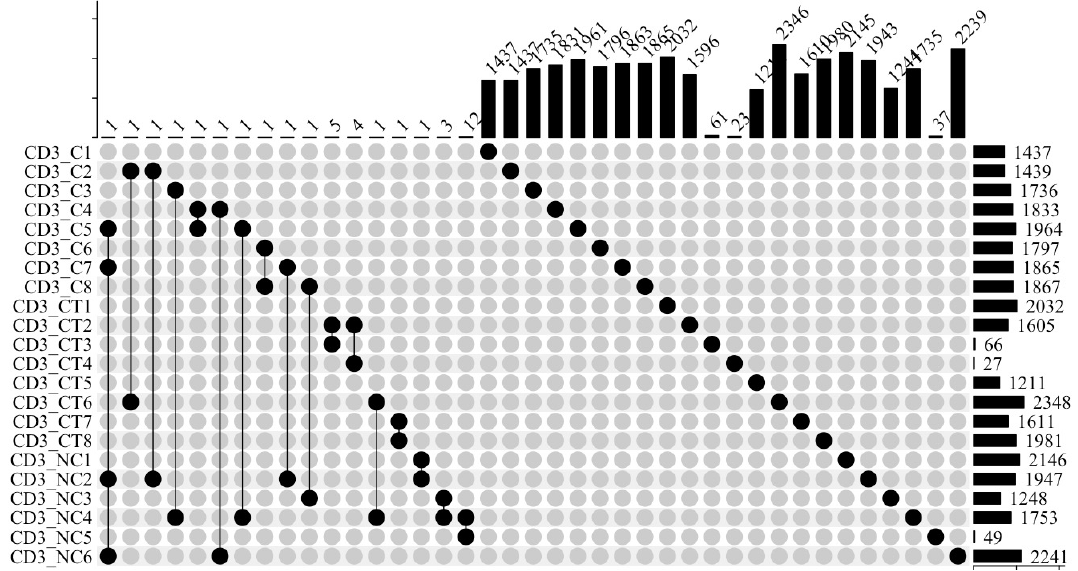

**Extended Figure R4. Colitis datasets showcase unique TCR sequence overlap.** The data is displayed as an UPSET plot. A dot indicate the presence of a unique clone. The line represents if there was any overlap across samples. (**top**) Bar graph represents the number of overlapping unique clone sequences that could be from multiple samples. (**right**) Bar graph representing the number of unique clones per sample. The uniqueness of the clones is based on the V(D)J arrangement of both chains and the unique CDR3 sequence of both chains. This plot includes both αβTCR and γδTCR lineages. C: colitis; NC: non-colitis; CT: normal controls.

We did not observe any clones that were specific to the cases experiencing colitis complication. Additionally, the repertoires of these individuals were also mostly private. Therefore, potential sequence similarity rather than exact matching may reveal TCR motifs more associated to a specific group (e.g., melanoma cases, no colitis etc.). To this end, TCR clustering with clusTCR2 was applied, resulting in the identification of 1607 alpha and 322 beta clusters (**Supplementary Table S6**), as well as 77 gamma and four delta clusters (**Supplementary Table S7**) that were present in one individual and a minimum of three clones.

Several clusters were found to be common among multiple individuals. These clusters included gamma clusters: TRGV9-TRGJP (n=21) and TRGV4-TRGJ2 (n=17) and alpha clusters TRAV1-2-TRAJ33 (n=18) and TRAV17-TRAJ54 (n=15). Despite their publicity, these patterns were not disease specific as they were observed across individuals with the different conditions. Indeed, the TRGV4-TRGJ2 is an expected cluster, as Vδ1γ4+ T cells are commonly observed in the colon (3). Similarly, TRGV9-TRGJP, commonly associated with the invariant Vδ2γ9+ T cells, is known to regularly interact with (E)-4-hydroxy-3-methyl-but-2-enyl pyrophosphate (HMBPP), a common microbial phosphoantigen (**Supplementary Table S7**) (4). The common alpha and beta clusters will be discussed in the context of the global analysis in **Results** **Section 3.4**. In conclusion, the identification of these gamma clusters is not unexpected in gut-derived T cells.

##### R3 Identification of disease specific phenotypes with semi-supervised annotation method

There is risk of not finding T-cell disease-specific patterns when interrogating a small number of cells or limited expression diversity due to experimental protocols (e.g., epitope-specific, tetramer sorting etc.). This is because T-cell functions are defined in contrast to other T-cell phenotypes, due to the minute variations that guide these differences. Consequently, identifying the specific signature may be missed due to expressional similarity or sparsity of the overall data present. This in turn limits our capacity to find the disease-specific/epitope-specific signature for any given TCR sequence. To overcome this challenge, we identified that combining all 12 datasets with 143 unique samples from 90 individuals allowed for identification of 49 distinct T cell phenotypes, including Th2 cells and DN populations that have been previously under-represented in studies. We examined the proportion of total CD8, CD4 and double negative (DN) T cells present. From the protein experience, specifically PBMCs, we would expect a bias of CD8>CD4> DN. Yet, this may not be the expected pattern when examining the transcriptome. The higher prevalence of DN is likely due to the poor coverage of the CD4+ marker. So researchers need to be cautions that the DN may be CD4+ T cells (**Extended Table R3**).

By combining the data, we could also interrogate stratification of the T cell population. The most common cell types (>5%) identified included (mean±standard error of the mean[SEM]; range): CD8ab+ Eff (25.0±2.6%; 12.3-38.5%), CD8ab+ Eff Tc1 (12.4±2.0%; 1 2.9-25.2%), CD8ab+ Eff Tc9 (5.9±2.0%; 0.4-24.0%), DN Naïve (7.0±1.2%; 0.4-14.5%) and DN FTH1 (5.5±1.2%; 0.4-14.2%).

There was substantial variation of the T cell sub-populations across the datasets. As expected, the tetramer (HPV epitope) sorted HNSCC was mostly CD8+ T cells (~97.0%) with less than 1% CD4+ T cells. Intriguingly, total naïve cell population were prevalent in breast cancer, COVID (lung tissue), T cell cancer (LGL), cervical cancer, Lewy body dementia (LBD) and acute myeloid leukemia (AML) (range 9.9% to 15.2%). Lastly there were considerable proportions of Tregs of both CD4+ and DN. The DN’s are likely to be CD4+ T cells, as the CD4 marker was poorly captured in 10x Genomics datasets. Tregs were highly prevalent (>8%) in breast cancer, esophageal cancer and melanoma, moderately present (5% to 8%) in colitis, LBD, anti-PD1 treated cancers, and lower percentage (<3%) in lung transplantation, HPV, COVID-19 lung tissue, LGL and AML.

##### R4 Public clones are likely related to common infections.

Given that we had access to 12 T-cell focused datasets, we next aimed to identify if there were any public clones across those studies, and if they could be present due to common histories (e.g., common viral infections). 53 of the ~250,000 unique clones were present in two or more studies (**Extended Table R4**). To further understand if these clones were common because of shared history (e.g., viral/bacterial exposure), we annotated these 53 αβTCRs with the DETECT tool for their epitope-specificity. 13 clones with a threshold of 0.2 (~0.99 probability) with high confidence of epitope specificity (**Extended Table R5**). This process identified nine TCRs with specificity for Human gammaherpesvirus 4 (Epstein–Barr virus [EBV]) for CLGGLLTMV, FLYALALLL, GLCTLVAML and YLRGRAYGL, three for Human betaherpesvirus 5 (Cytomegalovirus [CMV]) for NLVPMVATV and RPHERNGFTVL and one for Influenza A virus GILGLVFTL. The IMW DETECT could also identify eight clones that were likely MAIT cells as they were predicted to interact with MR1:5-OP-RU. The remaining 40 clones had unknown specificity.

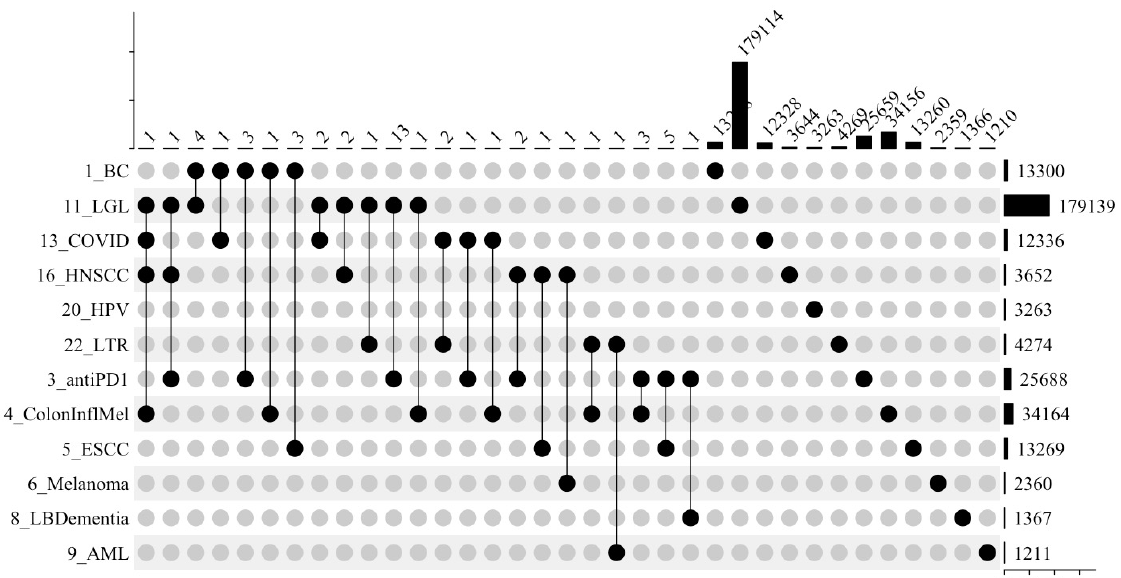

**Extended Figure R5. Overlap of clones across the 12 datasets.** (**top**) Bar graph represents the number of overlapping unique clone sequences that could be from multiple studies. (**right**) Bar graph representing the number of unique clones per study. The uniqueness of the clones is based on the V(D)J arrangement of both chains and the unique CDR3 sequence of both chains. This plot includes the αβTCR as the γδTCR were only present in the 4_ColonInflamMel data set.

##### R5 identification of MAIT cells with TCR-seq

MAIT cells traditionally have been identified based on the GEx alone annotation modelling that requires expression of TRAV1-2, KLRB1 and/or SLC4A10. To assess the enrichment of MAIT cells we first interrogated the standard GEx alone method using the unsupervised cluster-based annotation and FindMarker enrichment comparing the 25 clusters (resolution = 1). Only cluster 23 had significant enrichment of TRAV1-2 with TRBV6-2 expression and specifically to the Lewy body dementia (LBD) with 3.2% of their total cells (**Extended Figure R6A**). Next, we extracted the scaled expression of the TRAV1-2 gene to showcase that every cluster had this variable gene (**Extended Figure R6B**). Instead of using the GEx, we used the TCR-seq to identify the possible presence of MAIT cells using the TCR-seq based on the TRAV1-2 and J33/20/12. Interestingly, all data sets likely contained MAIT cell (GeoMean = 0.9%; SD±5.4%) with up to 19.3% of cells were likely MAIT cells in LBD (**Extended Figure R6C**). Overall, using the TCR-seq was better able to identify the presence of the MAIT population.

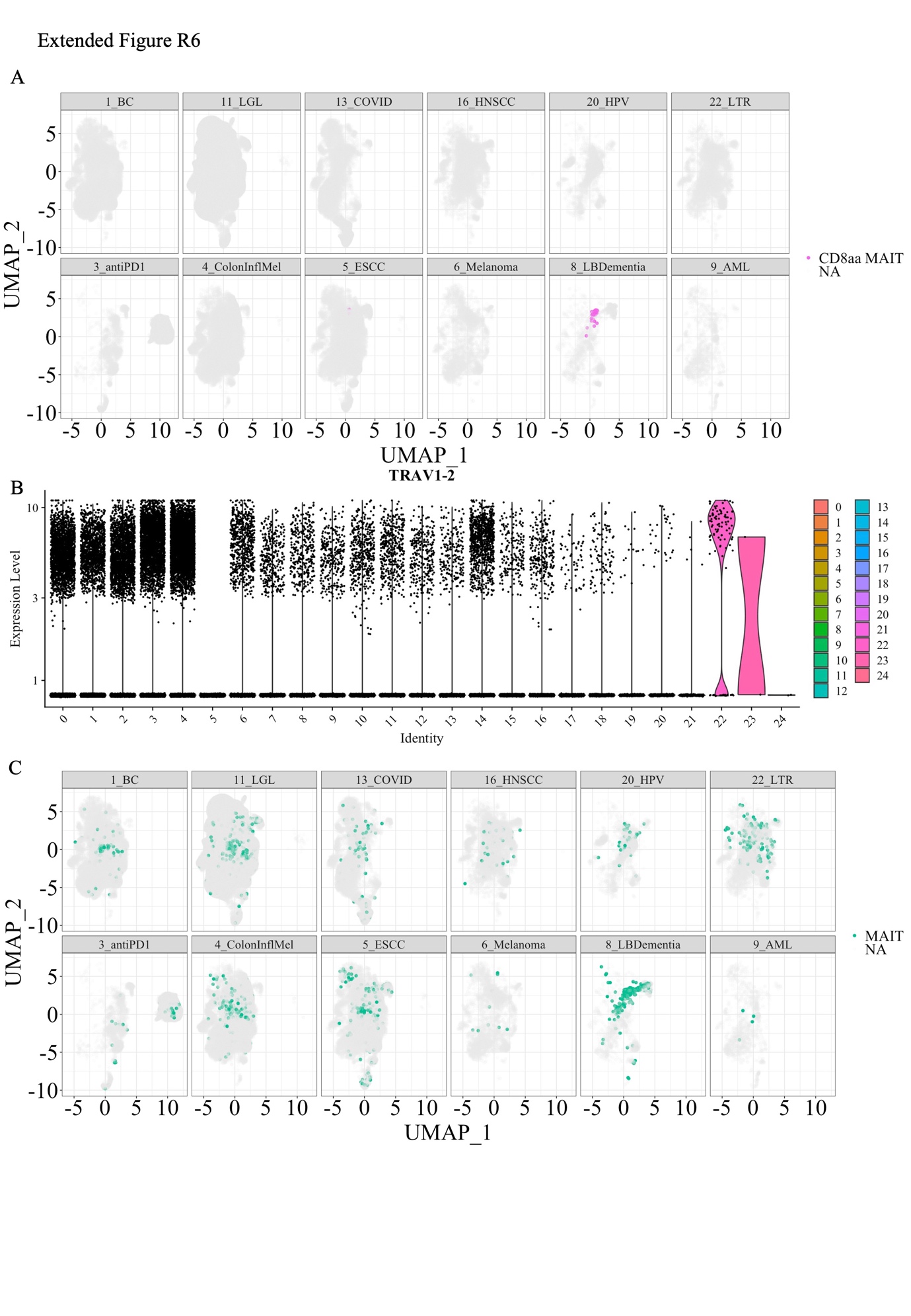

**Extended Figure R6. Identification of MAIT cells.** (**A**) UMAP plot of the Unsupervised clustering that was split by the different studies. This graph showcasing that only the lewy body (LB) dementia had a cluster of MAIT cells identified based on TRAV1-2 and KLRB1 expression. (**B**) Scaled gene expression of TRAV1-2 from the 12 studies was present in each of the clusters. (**C**) TCRseq MAIT annotation based on TRAV1-2 TRAJ33/12/20 were identified in every study.

### Extended Discussion

This extended discussion covers additional points related to single cell GEx analysis limitations as well as some dataset-specific insights.

An issue that not only plagues T cell datasets but is a wider problem is the consistency in annotating single cell datasets. Initially STEGO was developed as a tool for the dual analysis of TCR-GEx, but during the process we noted the pressing annotation issues related to defining T cell subpopulations. Not only did we note severe limitations, such as the neglection of the TCR in cell type definitions, the use of cell type markers was also markedly inconsistent across studies. Particularly, we identified several shortcomings of T cell annotations with regards to the unsupervised cluster and label transfer models. The unsupervised cluster process is appropriate for large population differences, using the TCR-seq we identified that there was large variability in the accuracy of CD4 vs CD8 T cells. The accuracy of this unsupervised clustering depends on which features maker up the PC’s to create the UMAP . IF they are not included as major different, their contribution is not readily incorporated into what makes up the UMAP. Additionally, we also identified that the TCR-seq had higher accuracy for finding the invariant T cell populations as well as highlighting γδTCR could be intermixed with the αβTCR. The unsupervised clustering does not take the TCR-specific inter account. As the label transfer models often are trained on the unsupervised annotation, they incur the same annotation inaccuracies as unsupervised clustering, as well as being limited by needing a highly similar dataset to be included for accuracy. Therefore, we do not recommend using unsupervised clustering for T cell annotations.

To ensure the annotation process was transparent and improved consistency, we chose the semi-automated gating-like annotation strategy with scGate (5) This process relies on defining the marker set (supervised aspect of the modelling), and levels (gating order) to find the population of interest. Testing ultimately identified having one marker per level and setting the threshold to 0.25 improved the overall accuracy of annotations that was checked with the CD8+ coverage. It was essential to also check the expression of each marker to ensure that they were consistent between the datasets and were not technical artifact. This strategy was applied to a combined dataset across all 149 samples with 500K cells (sorted on either CD3+ or CD45). The resulting data set contained 47 distinct annotated T cell phenotypes. Also, we chose to separate out some of the common states-based markers (senescence, immune checkpoint, cytotoxic, Th1-cytokine and cell cycling), as well as used the TCR-seq (e.g., invariant T cells), as these T cell features could co-occur with any of the 47 populations. To our knowledge, this is the first strategy that uses a joint GEx and TCR approach for annotation purposes. Also, for future use we uploaded the 500K TCR-GEx.rds object to Zenodo as a T cell atlas.

From the overall semi-supervised strategy, we identified disease specific features. One of the most notable findings was the changing prevalence of Tregs. Tregs are a T cell subset that induced self-tolerance, but in cancer setting can suppress anti-tumor immunity (6). This is because Tregs with IL-10 expression are causing the exhaustion in the CD4 and CD8+ T cell counter parts, which includes PD-1 expression (7). Here we identified many of the cancer types including breast cancer, esophageal cancer, head and neck cancer and melanoma, had a moderate to high levels over Tregs present. There was moderate over-representation of Tregs in the various anti-PD1 treatment cancer. There are varying response to anti-PD1 therapy with ~10% of cases with advanced gastric cancer having a serendipitous side-effect on increasing the Tregs population that lead to rapid cancer progression (8). So, in cancer settings inhibiting IL-10 rather than PD1 could be functionally interrogated to identify anti-IL-10 could be clinically beneficial in cancer settings. Overall, being able to compare many datasets enabled identification of disease-specific fluctuations in the T cell populations.

The original interrogations of the 12 studies could only use their own single cell data for the background. However, depending on the size of the included data this may lead to loss of signal. Here, we interrogated the lung transplantation recipient (LTR) experiencing acute rejection that interrogated the impact of glucocorticoid treatment. Comparing the expanded clone, which was mostly represented in the before therapy, there were too few clones (n=2) post therapy to determine the transcriptional changes. The overall profile comparing the expanded clone before treatment to the rest of the LTR showed that the cells were cytotoxic (GZMB) and HLA-DR genes (late activation). However, when this clone was enriched compared to the entire T cell Atlas, a distinct pattern emerged with several detoxification genes including the Metallothioneins MT1E, MT1F and MT1X. The increased expression of MT1 in regulatory T cells has been linked to lower IL-10 expression (9). Recent studies in mice that were IL-10+ had improved allograph tolerance compared to the IL-10 NULL population (10). We also noted that there were fewer Tregs in the LTR population. Potentially, the higher expression of metallothioneins may explain the lower prevalence of Tregs and needs to be a higher priority for experimentally validated in the context of acute cellular rejection in humans. Overall, using the entire T cell atlas enabled identification of transcriptional enrichment that was not identifiable in the original LTR due to the similarity of background issue. Overall, we recommend having adequate background data, and thus made the T cell Atlas available for this purpose.

Interrogating the GEx we could confirm that the HNSCC dataset was sorted, as they had very few CD4+ T cells present. Our next steps in re-interrogated the HNSCC dataset was how to be interrogate the tetramer sort that contained two HPV-epitopes KSA and QVD (11) . Focusing on TCRs that were both HPV-epitope specific that were also present in the PD-1 sort in primary tumor and metLN identified a possible HNSCC-epitope specific signature including RACK1 and LINC02446. RACK1 has been proposed as a regulator of T cell homeostasis [[36](bookmark://_ENREF_36)] or involved in activating CD4+ T cells [[37](bookmark://_ENREF_37)]. Here, we suggest it may be an indicator of activated CD8+ T cells as well. While little is known about the LINC02446 transcript, recent evidence implicates that it may be a promising therapeutic target in bladder cancer [[38](bookmark://_ENREF_38)]. Using the TCR-first framework we could identify and track the possible protective immune response generated by with the QVD and/or KSA epitopes in HLA-A*01:01+ individuals. Additionally, given the potential of these T cells to identify the cancers-HPV infected cells, engineering T cell (TCR-T) may aid in HLA-A*01:01+ non-responders. Collectively, this approach suggested alternative actionable features were identified that could involve targeting (e.g., RACK1 or LINC02446), preventative vaccination or T cell engineering.

In addition to resolving the annotation issue, we also determined how critical the TCR-seq was to identify if we need to include all lineages while interrogating single cell data. The current state of T cell-based research primarily centers around the αβTCR lineage, with limited attention given to γδTCR population. In the last 10 years γδ T cells have been identified to have diverse functions as innate- and adaptive-like (12), higher prevalence in mucosal membranes, and have emerging opportunity for novel therapies in cancer (reviewed in (13)). Single cell transcriptomics provides opportunity to overcome the current culturing limitations due to many of the epitopes remaining unknown, as this is no longer a barrier for interrogation. Nonetheless, γδ-T cells are rarely included in single cell studies due to not having the primers in the V(D)J kit (10x Genomics), as well as lacking optimizations in the downstream alignment processes (e.g. cell Ranger vdj alignment). Methods like TRUST4 may be used to partially recover γδ clones from scRNA-seq data, even without the inclusion of γδTCR primers, yet coverage remains limited at best (14). Of the 12 datasets only the colitis complication from melanoma therapy that had all four chains present, which a deliberate choice of the researchers due to gut being enriched for γδTCR populations (15). We noted a cluster of melanoma-specific TRGV4+ γδ T cells that expressed CD8ab+ T cell profile, and was indistinguishable from αβTCR counterparts. There is limited, but emerging evidence that γδ T cells can be tumor peptide-restricted with Vδ1γ5 with HLA-A*24:02 restriction (16), and Vγ4+ T cells with HLA-A*02:01 restriction to melanoma peptides (17). Given that the TRGV4+ cluster were CD8ab+, there is potential that they could have been peptide restricted. Overall, this interrogation highlights that we cannot assume all the CD8ab+ T cells are αβTCR, and currently the γδTCR and its contribution to disease settings is being missed due to lack of inclusion. We could elevate our T cell single cell data by including all four chains to reduce lineage biases (αβTCR vs γδTCR). Additionally, we could expand our focus based on both expansion, sequence similarity and TCR’s functional state to identify if they are likely to be peptide presenting or not.

The most ideal finding across individuals is the identification of a public clones. However, it is possible the public clones may not be disease specific, but due to other reasons. We compared the clones from all 12 datasets and identified 53 sequences. These TCR had limited expansion. The public clonotypes across disease setting were unlikely to be due to the specific pathology and were more likely due to a shared history. Approximately 1/3 of the sequences were predicted to known epitopes of EBV, CMV or MAIT related. The remaining 2/3 had effector phenotypes and may suggest that they would be epitope specific, but of an unknown origin. Overall, this analysis identified the rarity of finding public clones and recommendation using the prediction modelling to rule out if they are due to a common disease rather than the disease of interest.

To further understand the presence of non-disease specific patterns, we interrogated the sequence similarity and used our novel enrichment statistic. In addition to the TRAV1-2 TRAJ33 cluster that were likely MAIT cells, we identified that the TRAV3 and TRAV26-1 clusters also did not occur by random chance as it had more neighbors than expected for 51% and 41% of the cluster, respectively. Using the combination of hamming distance clustering, neighbor enrichment, GEx and IMW-DETECT was able to identify a novel TRAV3 cluster that could represent a shared disease history and ruled out a cluster that was not of interest.

**Extended references.**
